# A cysteine-rich domain of the *Cryptococcus neoformans* Cuf1 transcription factor is required for high copper stress sensing and fungal virulence

**DOI:** 10.1101/2024.12.13.628380

**Authors:** Corinna Probst, Chloe Insler, Catherine A. Denning-Jannace, Estefany Y Reyes, Jonathan L Messerschmidt, Lukas M. du Plooy, Charles Giamberardino, Connie B. Nichols, Yohannes Asfaw, Mari L Shinohara, Katherine J. Franz, J. Andrew Alspaugh

**Affiliations:** Department of Medicine, Duke University School of Medicine, Durham, NC, USA; Department of Molecular Genetics and Microbiology, Duke University School of Medicine, Durham, NC, USA; Department of Cell Biology, Duke University School of Medicine, Durham, NC, USA; Department of Integrative Immunobiology, Duke University School of Medicine, Durham, NC, USA; Department of Chemistry Duke University, Durham, NC, USA; Division of Laboratory Animal Resources Duke University, Durham, NC; Department of Chemistry and Biochemistry, University of North Carolina, Greensboro, USA; MRC Centre for Medical Mycology, University of Exeter, Exeter, UK

**Keywords:** nutritional immunity, metal homeostasis, fungal pathogenesis

## Abstract

The ability to sense, import and detoxify copper (Cu) has been shown to be crucial for microbial pathogens to survive within an infected host. Previous studies conducted with the opportunistic human fungal pathogen *Cryptococcus neoformans* (*Cn*) have revealed two extreme Cu environments encountered during infection: a high Cu environment within the lung and a low Cu environment within the brain. However, how *Cn* senses these different host Cu microenvironments, and the consequences of a blunted Cu stress adaptation for pathogenesis are not well understood. In contrast to ascomycete model fungi, the basidiomycete *Cn* has a single transcription factor (TF), *Cn*Cuf1, to regulate adaptive responses to both high- and low- Cu stress. Sequence comparison with other fungal Cu-responsive TFs identified three conserved cysteine (Cys)-rich motifs located within the *Cn*Cuf1 N-terminal domain, which were therefore predicted to play a role in Cu sensing. Mutation of these conserved Cys-rich motifs demonstrated that the 1^st^ Cys-rich motif is functionally relevant for *Cn*Cuf1 transcriptional activity during high Cu stress, while it is dispensable for low Cu stress adaptation. An inhalation model of murine infection showed that strains with defective high Cu stress regulation present a distinct and anatomically constrained pattern of yeast distribution within the infected lungs compared to a more widespread infection observed in lungs infected with the wild-type strain. Based on these findings, we hypothesize that Cuf1-driven high Cu responses modulate not absolute fitness but containment of *Cn* cells at the initial site of infection within the lung.

**Importance:** Copper is an essential micronutrient required for survival in all kingdoms of life as it is used as a catalytic cofactor for many essential processes in the cell. In turn, this reactivity of copper ions makes elevated levels of free copper toxic for the cell. This dual nature of copper – essential for life but toxic at elevated levels – is used by our innate immune system in a process called nutritional immunity to combat and kill invading pathogens. In this work we explore how the fungal human pathogen *Cryptococcus neoformans* senses high copper stress, a copper microenvironment encountered within the host lung. We identified a specific cysteine-rich motif within the copper responsive transcription factor Cuf1 to be essential for high copper stress sensing. Mutation of this motif led to an impaired high copper stress adaptation, which did not affect fitness of the yeast but did impact containment and distribution of yeast cells inside the host lung.

## Introduction

Copper (Cu) is an essential trace element in both microbes and mammalian cells, serving as a catalytic cofactor for a diverse range of cellular processes including iron (Fe) uptake and distribution; mitochondrial cytochrome oxidase activity; Cu, Zinc (Zn) superoxide dismutase catalysis; and melanin pigment production [1, 2]. Although Cu is required for life, elevated Cu levels are toxic due to its ability to induce uncontrolled redox cycling, causing the formation of toxic reactive oxygen species (ROS) and oxidative damage of DNA, proteins, and lipids [3, 4]. Free Cu ions were also shown to induce protein aggregation, triggering protein misfolding and precipitation [5, 6]. Furthermore, Cu has been shown to irreversibly displace Fe from Fe-S-cluster containing proteins, rendering these proteins nonfunctional [7, 8]. Therefore, organisms have evolved processes to sense and tightly regulate cellular Cu availability to maintain but not exceed the required cellular Cu quota [2, 4, 9-11].

Pathogenic microbes encounter unique extremes of Cu stress. Infected hosts often employ Cu-sequestering strategies known as “nutritional immunity” to limit Cu availability for invading pathogens [12-17]. In other sites, such as the macrophage phagolysosome, the host ATP7A Cu pump actively imports Cu to toxic concentrations to kill engulfed microorganisms [18]. In contrast, recent investigations demonstrated that brain resident microglia actually provide permissive conditions for the persistence of a microbial pathogen in the central nervous system (CNS) by supplying Cu in this otherwise nutritionally depleted site [13].

*Cryptococcus neoformans* (*Cn*) is an opportunistic fungal pathogen that causes life-threatening disease in immune compromised individuals [19, 20]. *Cn* is a free-living environmental fungus, frequently isolated from decaying plants, and soil contaminated with bird guano. Infectious spores or desiccated yeast cells are inhaled by mammalian hosts from the environment, resulting in an initial pulmonary infection [20]. In patients with weakened immune systems, including those with late-stage HIV infection, *Cn* disseminates through the bloodstream, crosses the blood-brain barrier, and colonizes the brain where it causes lethal infection [19, 20]. Previous studies have revealed that *Cn* encounters two extreme Cu environments during infection. During the initial infection within the lung, *Cn* senses a high Cu environment. In contrast, once the microorganism reaches the CNS, it senses low Cu availability [21, 22]. Thus, to maintain proper cellular Cu homeostasis, *Cn* needs to switch from Cu-detoxification during pulmonary infection to Cu-acquisition upon entering the CNS.

Several adaptive strategies for *Cn* Cu homeostasis have been described. To combat high Cu stress, *Cn* induces the Cu-detoxifying metallothionein genes *CMT1* and *CMT2*, the ROS detoxifying superoxide dismutase (*SOD*) 1 gene, as well as *ATM1*, a gene encoding a mitochondrial ABC transporter required to provide a sulfuric precursor for cytosolic Fe-S cluster biosynthesis [23]. These genes were previously shown to be required for *Cn* survival inside macrophages or in a mouse model of *cryptococcosis* [21, 24-26]. In contrast, within the Cu-restricted CNS environment, *Cn* induces the expression of its two high affinity copper importer genes, *CTR1* and *CTR4*, as well as the *CBI1* gene, encoding a Cu-binding protein required for efficient Cu acquisition [22, 27, 28]. Additionally, recent investigations defined a tightly regulated cellular switch between the Cu/Zn-dependent Sod1 to the Cu-independent/manganese (Mn)-dependent Sod2 during Cu limiting conditions. This switch allows the fungal cell to prioritize Cu for use in other essential Cu-dependent cellular processes when labile Cu is limited without disturbing intracellular ROS detoxification [11].

Cu sensing and regulation has been well studied in ascomycete model fungi, of which most employ two distinct transcription factors (TFs) to control responses to high and low Cu stress [2]. For example, the *Saccharomyces cerevisiae* (*Sc*) Ace1 transcription factor is activated in a high Cu environment to regulate gene expression to protect the yeast from toxic Cu levels [29, 30], and the *Sc* Mac1 transcription factor is activated by low Cu environments to favor Cu acquisition [31]. In contrast, the basidiomycete *Cn* maintains a single Cu-responsive transcription factor, Cuf1, that controls both high and low Cu stress-responsive genes [23]. Deletion of *CUF1* renders *Cn* less virulent in an intravenous mouse model of *cryptococcosis* [32], further underscoring the importance of Cu homeostasis in *Cn* pathogenesis and suggesting the potential of *Cn*Cuf1 as a fungal-specific intervention target to disrupt pathogen-driven adaptations to the host nutritional environment during infection. However, the actual mechanism of high and low Cu gene regulation by Cuf1 in response to various Cu stresses is not well understood and has yet to be elucidated.

*Cn*Cuf1 was named and identified based on sequence homologies with the Cu-responsive TF found in *Schizosaccharomyces pombe* (*Sp*), with which it shares the highest sequence homology [32, 33]. However, unlike *Sp*Cuf1, which only localizes to the nucleus to direct gene expression of its target genes during Cu limitation, *Cn*Cuf1 is bound to and directs expression of different subsets of target genes during either low or high Cu stress [23, 34]. Despite *Cn*Cuf1’s distinct function as dual Cu-responsive TF, it shares conserved, known functional relevant motifs with the previously studied ascomycete Cu-responsive TFs. To further define the molecular mechanisms driving the unique dual activity of this Cu-responsive TF, we mutated these potential functional protein motifs in *Cn*Cuf1 and tested the effects of these mutations on fungal physiology and stress responses. We also explored the consequences of *Cn*Cuf1 mutations targeting the *Cn*Cuf1 high Cu stress response for pathogenesis-relevant cupro-enzyme activity and its effects on pathogenesis within the mouse lung environment using an inhalational mouse model of *cryptococcosis*.

## RESULTS

### Mutation of the Ace1-like Cys-rich motif in the Cuf1 transcription factor renders the fungus susceptible to high Cu stress

*Cn*Cuf1 is the central Cu-responsive transcription factor known to regulate high and low Cu genes in *Cn* (Fig 1). Sequence comparison between *Cn*Cuf1 and other fungal Cu-responsive transcription factors identified the highest homology within the first 39 amino acids (aa) of the N-terminal domain [23]. This region includes a Zn-binding motif and the highly conserved, positively charged ‘KGRP’ motif [29, 31, 35-38]. Previous mutational studies in *Cn*Cuf1 have already demonstrated that the KGRP motif is required for promoter binding of both high and low Cu-regulated genes [23] (Fig 1).

**Figure 1:**
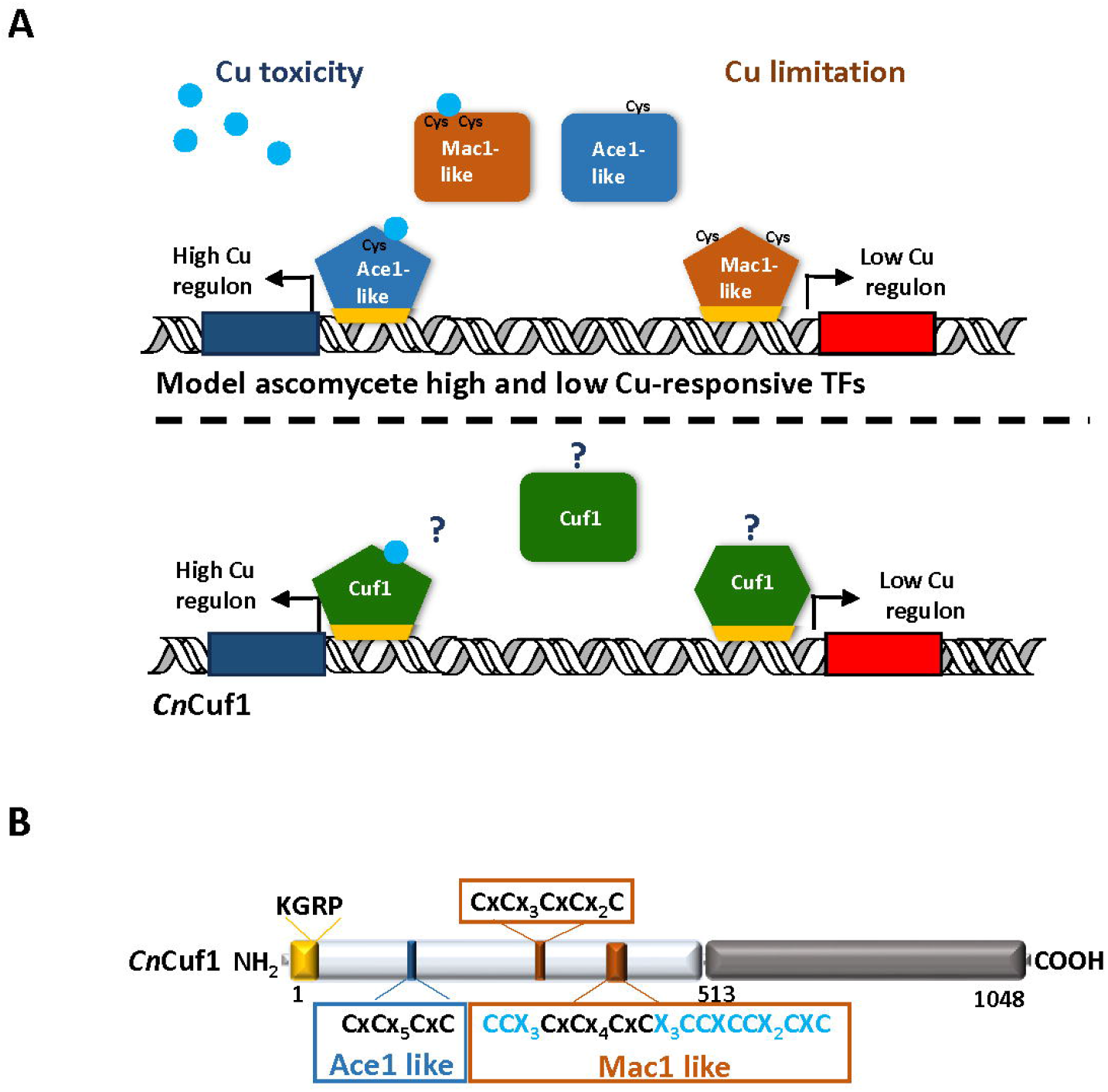
Comparison of single versus dual fungal Cu-sensing transcription factors. **(A)** Mode of action of single Cu-sensing TFs found in model ascomycetes versus the dual Cu-sensing TF in *Cn*. Both systems use a highly conserved DNA binding motif to bind target promoter sequences (highlighted in yellow). Cu binding to Cys-rich motifs encoded by single Cu-sensing TFs results in confirmational changes, which either inhibit transcriptional activity of low Cu-responsive Mac1-like TFs (orange) or activate transcriptional activity of high Cu-responsive Ace1-like TFs (blue). In contrast, *Cn* uses only one Cu-responsive TF, *Cn*Cuf1 (green), which must remain transcriptionally active during low and high Cu stress but is directed to distinct promoter regions based on variance of cellular Cu levels. Motifs involved in Cu-sensing and regulating *Cn*Cuf1 transcriptional activity are yet unknown. **(B)** Schematic representation of the *Cn*Cuf1 domain structure and conserved motifs. The first and last residues of each domain are indicated. The N-terminal domain is shown in light grey; the C-terminal domain is shown in dark grey. The N-terminal DNA binding motif, which contains the conserved KGRP DNA binding motif, is highlighted in yellow. The Ace1-like cysteine-rich region is indicated via a blue box, and the Mac1-like regions are indicated by an orange box. The sequence of the identified cysteines (Cys, C) is indicated for each region; the *Cn*Cuf1-unique Cys-rich sequence located adjacent to the 2^nd^ Mac1-like Cys-rich motif is highlighted in turquoise.

Following the *Cn*Cuf1 DNA-binding domain, the sequence similarity decreases considerably, except for three cysteine (Cys)-rich motifs, which were previously demonstrated in ascomycete high and low Cu-responsive TFs to be essential for regulation of transcriptional activity; Cu binding to these Cys-rich motifs acts as an ON-OFF switch either activating (Ace1-like transcription factors) or deactivating (Mac1-like transcription factors and *Sp*Cuf1) transcriptional activity [29, 34-36, 39] (Fig 1). The first Cys-rich motif encoded by *CUF1* is formed by an 11 aa motif, containing 4 cysteines, which are highly conserved among Cuf1- and Ace1-like transcription factors [23] (Fig 1B). To elucidate the function of the Ace1-like Cys-rich motif on *Cn*Cuf1 transcriptional activity, we mutated all 4 conserved Cys located in the Ace1-like Cys rich motif to alanine (Ala, Cuf1-FLAG ΔAce1, Supp. Fig 1A). The 2^nd^ and 3rd Cys-rich motif encoded by *CUF1* are formed by a 12 aa motif containing 5 Cys (2^nd^ Cys-rich motif) and a 10 aa motif containing 4 cysteines (3^rd^ Cys-rich motif), which are highly conserved among Cuf1- and Mac1-like transcription factors [23] (Fig 1B). Additionally, we identified a sequence of 7 cysteines adjacent to the Mac1-like 3^rd^ Cys-rich motif, which are unique to *Cn*Cuf1 (Fig 1B). To elucidate the function of the Mac1-like Cys-rich motif on *Cn*Cuf1 transcriptional activity, we mutated all 9 conserved cysteines located in the two Mac-like Cys rich motifs encoded by *Cn*Cuf1, as well as the 7 adjacent cysteine residues to alanine (Cuf1-FLAG ΔMac1, Supp. Fig 1A). Alleles encoding FLAG-tagged versions of the Cuf1 wild type (WT) protein, the KGRP-mutant protein, the Cuf1 ΔAce1 mutant and the Cuf1 ΔMac1 mutant proteins were incorporated into the *cuf1Δ* mutant strain at the safe haven 1 site [40], all expressed by the *CUF1* promoter.

To explore the effects of the introduced mutations on Cuf1-FLAG protein levels at high and low Cu stress, we conditioned the *CUF1*-FLAG-expressing strains for 3 h in either SC medium + 1 mM CuSO_4_ (= high Cu stress condition) or SC medium + 1 mM of the extracellular Cu chelator bathocuproinedisulfonic acid (BCS,= low Cu stress condition) at 30°C. Western blot analysis using an anti-FLAG-HRP conjugate demonstrated no differences in Cuf1-FLAG protein levels in response to high or low Cu stress between the WT and any of the Cuf1 mutant proteins, indicating that the introduced site mutations do not affect Cuf1 protein levels (Fig 2A).

**Figure 2:**
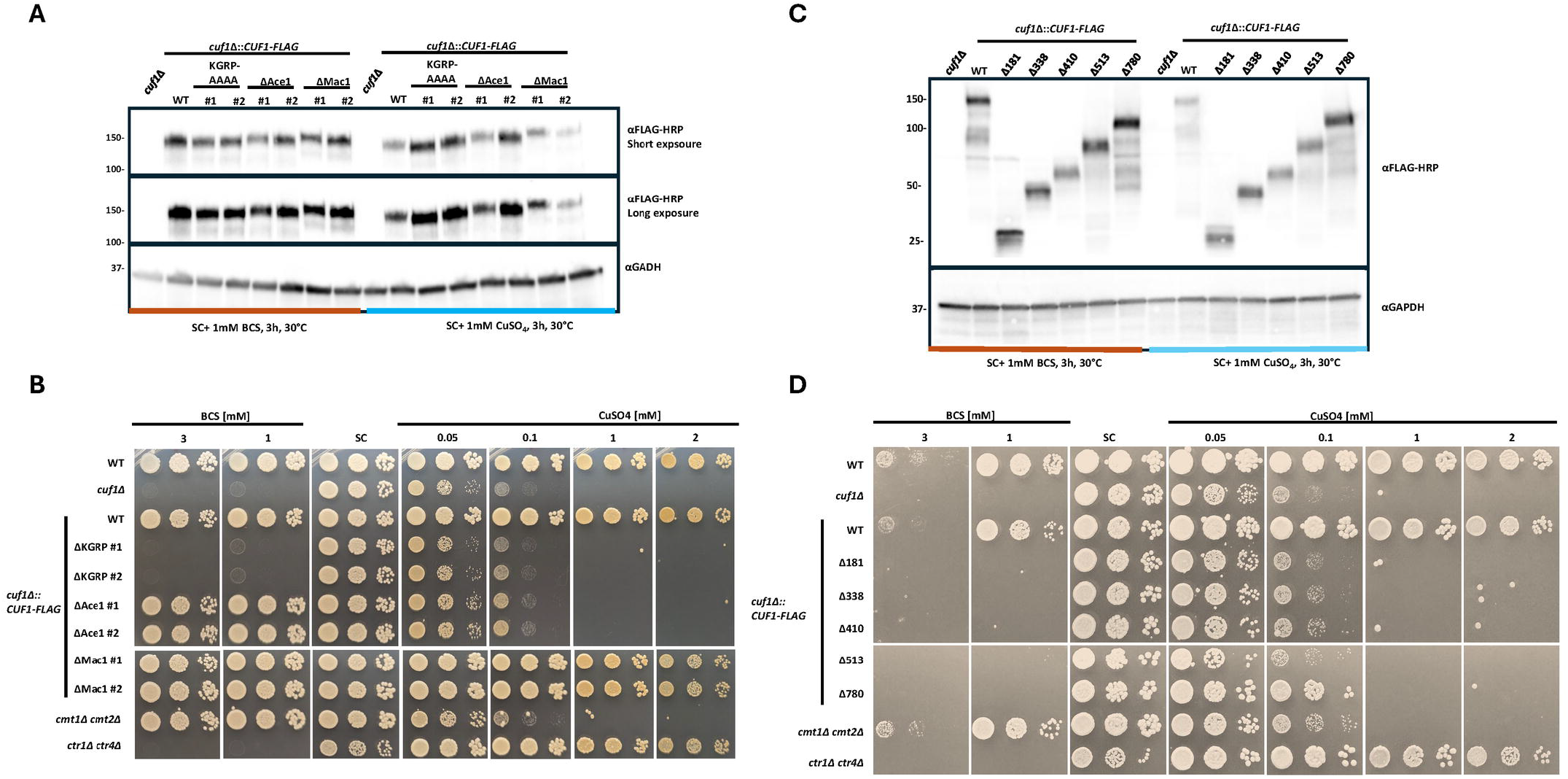
The Cuf1-FLAG ΔAce1 mutant protein is produced to WT levels but fails to restore growth during high Cu stress. **(A)** Cuf1-FLAG WT, Cuf1-FLAG KGRP-AAAA, Cuf1-FLAG ΔAce1 and Cuf1-FLAG ΔMac1 mutant protein expression assessed by western blot analysis. Indicated *CUF1-FLAG* expressing strains were incubated in SC medium supplemented with either 1 mM CuSO_4_ or 1 mM BCS at a starting OD_600_ of 0.5 for 3 h at 30°C. A TCA based protein extraction was performed. 25 µL of crude TCA extract was analyzed per lane with the α FLAG-HRP conjugate. The α GAPDH (anti-GAPDH) antibody was used as a loading control. **(B)** Growth analysis in the presence of high and low Cu stress. Five-fold serial dilutions of cell suspensions for each strain were spotted onto SC medium agar plates. To induce low Cu stress, the SC medium was supplemented with 0.5 or 1 mM BCS. To induce varying degrees of high Cu stress, the SC medium was supplemented with either 0.02, 0.05, 0.1, 0.25 or 1 mM CuSO_4_. Plates were incubated at 30°C for 3 days. Two independent strains of each *CUF1-FLAG* mutant were analyzed. As Cu stress controls, the Cu import mutant *ctr1*Δ *ctr4*Δ and the Cu detoxification mutant *cmt1*Δ *cmt2*Δ were included in the growth analysis. **(C)** Cuf1-FLAG WT, Cuf1-FLAG Δ181, Cuf1-FLAG Δ338, Cuf1-FLAG Δ410, Cuf1-FLAG Δ513 and Cuf1-FLAG Δ780 protein fragment expression assessed by western blot analysis. Indicated *CUF1-FLAG* expressing strains were incubated in SC medium supplemented with either 1 mM CuSO_4_ or 1 mM BCS at a starting OD_600_ of 0.5 for 3 h at 30°C. Protein extraction and western blot was performed as described in (A). **(D)** Growth analysis in the presence of high and low Cu stress. Five-fold serial dilutions of cell suspensions for each strain were spotted onto SC medium agar plates. Indicated strains were spotted on to Cu stress plates as described in (B).

To determine the physiological effect of the introduced site mutations on regulation of Cu homeostasis during Cu stress, we assessed the ability of each allele to complement the *cuf1Δ* growth phenotypes in high and low Cu stress environments. In all stress conditions, the *CUF1-FLAG* WT allele fully restored growth of the *cuf1Δ* mutant strain, demonstrating that the FLAG-tag did not compromise Cuf1 protein function (Fig 2B). In contrast, the KGRP-mutant protein, which was previously demonstrated to lack *Cn*Cuf1-promoter binding capability, predictably did not complement the growth phenotypes of a *cuf1Δ* mutant strain [23]. Consistent with the postulated function of the Ace1-like Cys-rich motif for regulating high Cu stress responses, a *CUF1*-*FLAG* ΔAce1 mutant allele failed to complement the *cuf1Δ* growth defect in a high Cu environment, however it fully complemented the *cuf1Δ* low Cu growth phenotype (Fig 2B). This finding suggests a specific role for the Ace1-like Cys rich motif for regulating transcriptional activity in response to high Cu stress. In contrast to the assumed function of the two identified Mac1-like Cys-rich motifs for low Cu stress sensing, the Cuf1-Flag ΔMac1 mutant allele fully complemented the *cuf1Δ* growth defects in presence of both high and low Cu stress. This finding suggests that *Cn*Cuf1 senses and regulates low Cu stress independent from the Mac1-like Cys-rich motifs (Fig 2B).

### The *Cn*Cuf1 C-terminal unstructured domain is required for regulating *Cn*Cuf1transcriptional activity

In addition to the N-terminally located conserved DNA binding domain and the Cys-rich motifs, Cn*CUF1* encodes a C-terminal unstructured domain of unknown function, which is unique to *Cn*Cuf1 and not found in any of the studied ascomycete Cu-responsive TFs [23]. The biological function of this extended unstructured C-terminus, and its role for Cu-dependent regulation of *Cn*Cuf1 activity, are unknown. To better understand and define regions required for high and low Cu stress adaptation, we performed a sequential C-terminal truncation analysis of the *Cn*Cuf1 transcription factor using the *CUF1-FLAG* WT allele-encoding plasmid (Supp. Fig 1B). Alleles encoding the FLAG-tagged versions of the generated *CUF1-FLAG* truncations were incorporated into the *cuf1Δ* mutant strain at the safe haven 1 genomic site [40], all expressed by the *CUF1* promoter.

To explore the effects of the introduced truncations on Cuf1-FLAG protein levels at high and low Cu stress, we conditioned the *CUF1*-*FLAG* truncation-expressing strains as described above and performed western blot analysis using an anti-FLAG-HRP conjugate. All generated *CUF1-FLAG* truncations were expressed at similar protein levels independent from the Cu stress condition (Fig 2C).

Like the generated *CUF1* Cys-site mutant strains, we assessed the ability of each *CUF1-FLAG* fragment to complement the *cuf1Δ* growth phenotype in high and low Cu stress environments. Surprisingly, none of the generated *CUF1-FLAG* fragments fully complemented the *cuf1Δ* mutant phenotypes in either Cu stress condition (Fig 2D). Only the largest generated *CUF1-FLAG* fragment (Δ780), which still encodes for a portion of the unstructured C-terminus unique to *Cn*Cuf1, showed a partial growth restoration of the *cuf1Δ* high Cu stress growth defect (Fig 2D, 0.1mM CuSO_4_). This finding suggests an important role of the unique C-terminal region for regulation of *Cn*Cuf1 transcriptional activity.

### The Ace1-like Cys-rich motif is required for Cuf1 to specifically regulate high Cu-responsive genes

To correlate the observed growth phenotypes of the *CUF1-FLAG* Cys-rich motif mutant strains with *Cn*Cuf1 transcriptional activity, we used a dual fluorescent Cu sensor strain to simultaneously measure the activation of the low Cu-regulated *CTR4* gene, encoding for a high affinity Cu importer, as well as the high Cu-regulated *CMT1* gene, encoding for a Cu detoxifying metallothionein. To create the Cu sensor strain, a *CTR4* promoter (p*CTR4*)*-*driven *mCLOVER* gene and an *CMT1* promoter (p*CMT1*)-driven *mRUBY* gene were inserted into the safe haven 2 site of the *Cn* genome, a “neutral” genomic locus into which new allele introduction does not result in alterations of physiology or virulence [41](Supp. Fig 2A). Doing so resulted in a strain expressing green fluorescence during Cu limitation (induction of p*CTR4* driven *mCLOVER* expression) and expressing red fluorescence during excess Cu (induction of p*CMT1*-driven *mRUBY* expression, Supp. Fig 2B-C). To measure the effects of the *CnCUF1* Cys-rich motif mutant alleles on *Cn*Cuf1 transcriptional activity, we introduced the green- and red-fluorescent Cu sensor alleles into the WT and the *cuf1Δ* mutant; as well as the *cuf1Δ* mutant strain expressing either the *CUF1-FLAG WT* allele, *CUF1-FLAG KGRP-AAAA* mutant allele or the *CUF1-FLAG* Cys-rich motif mutant alleles. All strains were cultured for 24 h in SC medium supplemented with either 50 µM CuSO_4_ (=excess Cu) or 50 µM BCS (=low Cu) at 30°C prior to protein extraction and western blot analysis of mClover and mRuby protein levels (Fig 3A).

**Figure 3:**
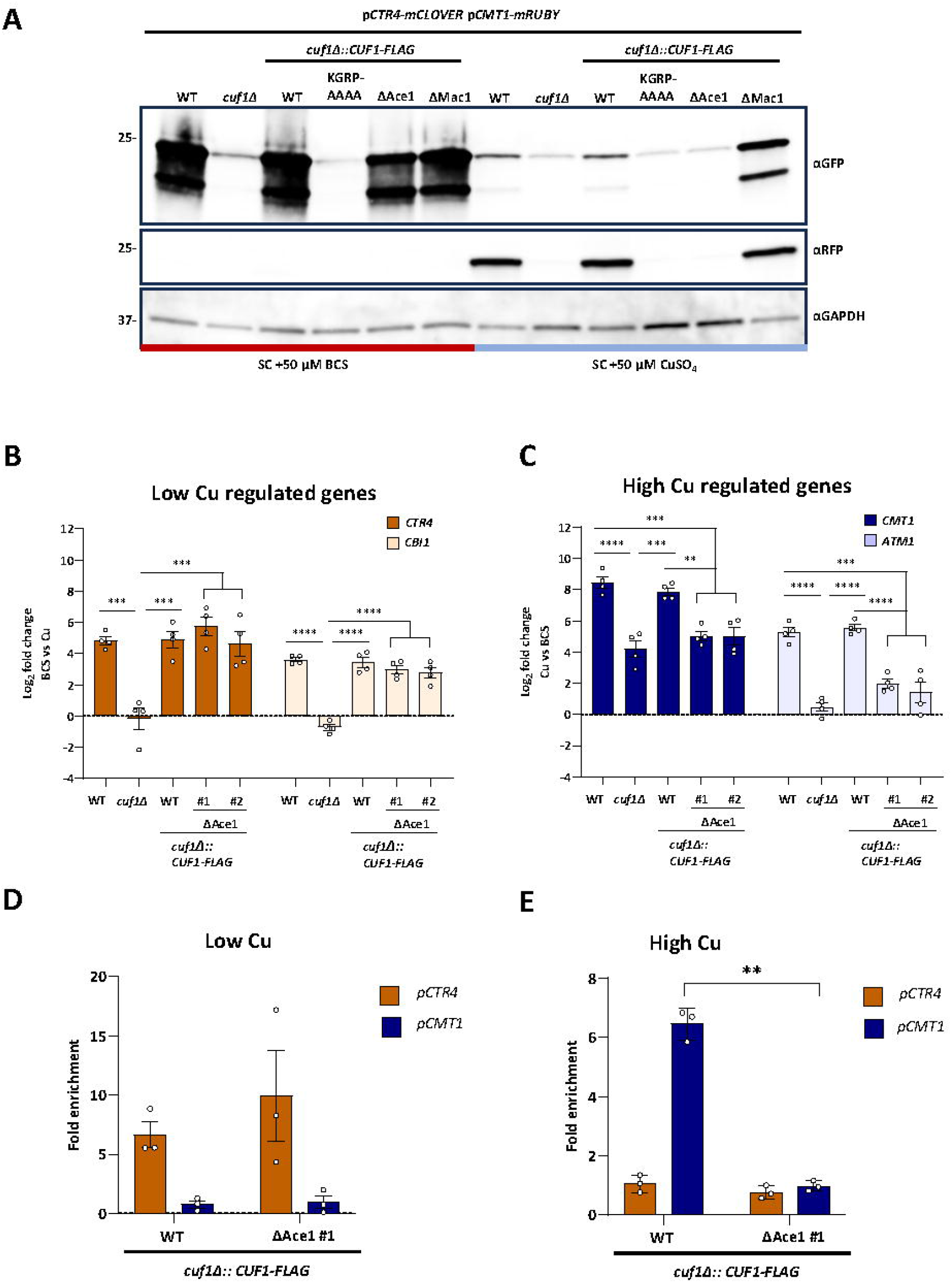
Mutation of the Ace1-like Cys-rich motif renders Cuf1 unable to bind to and activate high Cu regulated genes. **(A)** p*CTR4*-m*CLOVER* and p*CMT1*-m*RUBY* expression profiles of *CUF1-FLAG* WT and site mutant expressing strains. Indicated strains were inoculated to OD_600_ of 0.1 into SC + 50 µM BCS or SC + 50 µM CuSO_4_ and grown for 24 h at 30°C. A TCA based protein extraction was performed and 20 µg total protein was analyzed per lane. mClover was detected using a monoclonal anti-GFP antibody (mouse, αGFP) and mRuby was detected using a polyclonal anti-RFP antibody (rabbit, αRFP). An anti-GAPDH antibody (rabbit, αGAPDH) was used as a loading control. **(B-C)** Transcript abundance of the low Cu-regulated *CTR4* and *CBI1* genes (B) and the high Cu-regulated *CMT1* and *AMT1* genes (C). Indicated strains were diluted to OD_600_ of 0.5 and incubated for 2 h at 30°C in SC medium. Cu stress was induced by the addition of either 1 mM BCS or 1 mM CuSO_4_ and cultures were incubated for 3 h at 30°C. Transcript abundance for each indicated gene was assessed by quantitative real-time PCR (qPCR), and the log_2_ fold change of transcripts was calculated using the ΔΔ C_T_ method in low Cu stress versus high Cu stress (B) or high Cu stress versus low Cu stress (C). Transcript levels were normalized to *GAPDH*. Shown is the mean +/-SEM of 4 biological replicates. Statistical analysis was performed using one-way ANOVA. **(D-E)** Cuf1-FLAG promoter binding at low Cu stress (D) or high Cu stress (E). Indicated strains were diluted to OD_600_ of 0.25 and incubated for 2 h at 30°C in SC medium. Cu stress was induced by the addition of either 1 mM BCS or 1 mM CuSO_4_, and cultures were incubated for 2.5 h at 30°C prior to crosslinking. Promoter binding to the target promoters was analyzed via (ChIP-PCR). Fold enrichment was calculated by comparison of target promoter abundance in the FLAG-tagged strains to the promoter abundance in the non-FLAG tagged, WT strain. The mean +/-SEM of 3 biological replicates is shown. A two-tailed t-test was performed as statistical analysis.

Consistent with previously published transcript analysis of Cu- and Cuf1-dependent genes [23], the WT Cu-sensor strain showed robust activation of p*CTR4*-driven *mCLOVER* expression when grown in Cu-limiting conditions (SC+ 50 µM BCS) and robust activation of p*CMT1*-driven *mRUBY* expression during Cu excess (SC +50 µM CuSO_4_). As predicted by the established activity of *Cn*Cuf1 to control the expression of low and high Cu-responsive genes, deletion of the *CUF1* gene resulted in a failure of Cu-dependent gene activation as measured by the absence of mClover and mRuby protein expression (Fig 3A). Introduction of the *CUF1-FLAG* WT allele, which fully complemented the Cu-stress growth defects of the *cuf1Δ* mutant strain, also fully restored the Cu-dependent mClover and mRuby protein levels to WT-like levels. In contrast, expression of the *CUF1-FLAG* KGRP-AAAA DNA binding mutant, which failed to complement the *cuf1Δ* mutant growth phenotypes, showed similarly absent *mCLOVER* and *mRUBY* expression as the *cuf1Δ* mutant strain.

In line with their ability to complement the *cuf1Δ* low Cu stress growth phenotype, neither Cys-rich motif mutations (ΔAce1 or ΔMac1) affected *mCLOVER* expression levels during Cu limitation (Fig 3A). However, a slightly higher m*CLOVER* expression was observed for the *CUF1*-FLAG ΔMac1 strain when grown in Cu excess, suggesting a potential role for the Mac1-like Cys-rich motifs to facilitate complete repression of low Cu-regulated genes, such as *CTR4*, when Cu is in excess. No effect on *mRUBY* expression was observed in the *CUF1*-FLAG ΔMac1 expressing strain at either Cu conditions, suggesting that these Mac1-like regions are not involved in the activation of genes induced in high Cu environments. In contrast, mutation of the Ace1-like Cys-rich motif fully inhibited the high Cu-driven activation of *mRUBY* expression. Hence, the previously observed growth defect of the *CUF1*-FLAG ΔAce1 expressing strain in presence of high Cu stress is likely due to failed activation of genes required for Cu detoxification. Together, the Cu sensor expression data as well as the Cu stress growth data suggests that mutation of the Ace1-like Cys-rich motif results in an uncoupling of *Cn*Cuf1-directed high Cu and low Cu stress adaptation, with the high Cu stress adaptation being completely blunted while the low Cu stress adaptation seems to be fully functional.

To further validate the observed gene expression phenotype of the *CUF1-FLAG*-ΔAce1 mutant strain, we directly measured transcript levels for established high and low Cu-regulated genes using quantitative reverse transcriptase PCR (qRT-PCR) in the WT strain, the *cuf1*Δ mutant strain, as well as the *cuf1*Δ mutant strain expressing either the *CUF1-FLAG WT* allele or the *CUF1-FLAG* ΔAce1 allele. As a measure of high Cu stress transcriptional responses, we measured transcript abundance of the *CMT1* gene, whose promoter was used in the Cu sensor strain, and the *ATM1* gene, encoding for a mitochondrial inner membrane ABC transporter involved in cytosolic Fe-S assembly [21, 24]. To assess responses to low Cu stress, we measured the transcript abundance of the *CTR4* gene, whose promoter was used in the Cu sensor strain, and the *CBI1* gene, encoding the copper binding and release protein 1 [22, 23, 28].

The above indicated strains were incubated for 3 h in SC medium supplemented with either 1 mM CuSO_4_ (=high Cu stress condition) or 1 mM BCS (=low Cu stress condition) at 30°C prior to RNA extraction. Transcript levels of the Cu-response genes for each strain in each condition were determined by qRT-PCR. Consistent with prior results, the expression of the *CMT1* and *ATM1* genes was induced in the WT strain under high Cu conditions, and the *CTR4* and *CBI1* genes were induced in low Cu conditions (Fig 3B-C) [23]. In contrast, the transcript levels of both high Cu and low Cu genes were not induced in either condition in the *cuf1*Δ mutant. The *CUF1-FLAG* WT strain fully restored WT levels of Cu-regulated gene expression in both conditions. However, consistent with the observed Cu stress growth phenotypes and the observed mClover and mRuby protein expression profile, mutation of the Ace1-like Cys-rich motif was associated with defective transcriptional activation of high Cu-regulated genes (*CMT1* and *ATM1*, Fig 3C), while fully restoring WT transcript levels of genes involved in low Cu stress response (*CTR4* and *CBI1*) (Fig 3B).

### The Ace1-like Cys-rich motif is required for Cuf1 binding to high Cu-responsive gene promoters

The Cuf1 transcription factor was previously demonstrated to directly bind the promoters of these high- and low-Cu reporter genes [23]. We used chromatin immunoprecipitation followed by quantitative PCR analysis (ChIP-PCR) to determine whether the observed differential transcript levels in each strain were associated with changes in Cuf1 promoter binding. We specifically chose oligonucleotides corresponding to the *CTR4* and *CMT1* promoters as indicators of low- and high-Cu stress gene promoter binding, respectively. To do so, we incubated the strains in the identical high- and low-Cu stress conditions as above prior to protein-DNA crosslinking and chromatin immunoprecipitation. Enrichment of the *CMT1* or *CTR4* promoter fragments in each sample was calculated relative to the WT strain and was further normalized against the *TUB1* tubulin gene as a non-Cuf1 target gene. Consistent with previous studies, we observed a significant enrichment of the *CMT1* promoter, but not the *CTR4* promoter, in the Cuf1-FLAG WT sample conditioned in high Cu stress, and enrichment of the *CTR4* promoter sequence during low Cu stress [23](Fig 3D-E). In contrast, no enrichment of the *CMT1* promoter was observed by ChIP-PCR in the *cuf1*Δ + *CUF1-FLAG* ΔAce1 strain conditioned in high Cu stress, whereas the *CTR4* promoter was enriched in this strain to similar levels as in the Cuf1-FLAG WT allele expressing strain (Fig 3D-E). Taken together, these data demonstrated that the Ace1-like Cys-rich motif is required for Cuf1 binding to and regulation of high Cu-responsive promoters while it does not impact binding or regulation of low Cu-responsive promoters.

Together, these transcript and promoter binding data demonstrate that the Ace1-like Cys-rich motif is required for the *Cn*Cuf1 transcription factor to transcriptionally activate high Cu-responsive genes but not low Cu-responsive genes. This activity likely explains the defective cellular adaptation to high Cu stress for strains with mutations of this site.

### The Ace1-like Cys-rich motif is required for proper regulation of ROS detoxification and melanization in high Cu environments

*Cn* encounters both low- and high-Cu microenvironments in the infected host [22]. A full loss-of-function mutation of the *CUF1* gene ultimately affects both high and low Cu stress adaptation, which complicates the analysis of high Cu or low Cu stress responses for *Cn* pathogenesis. Our mutational analysis identified the *CUF1-FLAG*-ΔAce1 mutant strain as a candidate strain to specifically analyze the role of high Cu stress adaptation for *Cn* pathogenesis. First, we assessed the role of Cuf1 and the Ace1-like Cys-rich motif for the activity of cupro-enzymes required for *Cn* stress resistance and survival within specific microenvironments of the human host.

#### Laccase 1 and melanin production

The *Cn*Lac1 protein is a Cu-containing diphenol oxidase/laccase involved in formation of melanin, a cell wall-associated pigment that protects the fungus from host-derived oxidative and nitrosative stress as well as from antifungal drugs [42]. The process of melanization is tightly linked to Cu homeostasis and Cuf1 regulation in multiple ways. First, melanin production requires cellular Cu as Lac1 oxidase activity is dependent on Cu as a catalytic cofactor [43]. Second, *LAC1* gene expression is controlled in a Cuf1-dependent manner [23, 27, 44, 45](Fig 4A). To analyze the effects of the Ace1-like Cys site mutation on melanin production, we incubated the WT strain, the *cuf1*Δ mutant, and the *cuf1*Δ strain expressing either *CUF1-FLAG* WT or the ΔAce1-like Cys site mutant allele on L-DOPA medium supplemented with either 10 µM BCS (to slightly restrict Cu) or 10 μM CuSO_4_ (to provide excess Cu). As controls, we included the completely melanin-defective *lac1*Δ mutant strain and the *cbi1*Δ *ctr4*Δ double mutant strain that is impaired in copper import [28, 43]. For each strain, relative *LAC1* transcript levels were measured after 3 h of incubation in liquid L-DOPA medium similarly supplemented with either 10 µM BCS or 10 μM CuSO_4_.

**Figure 4:**
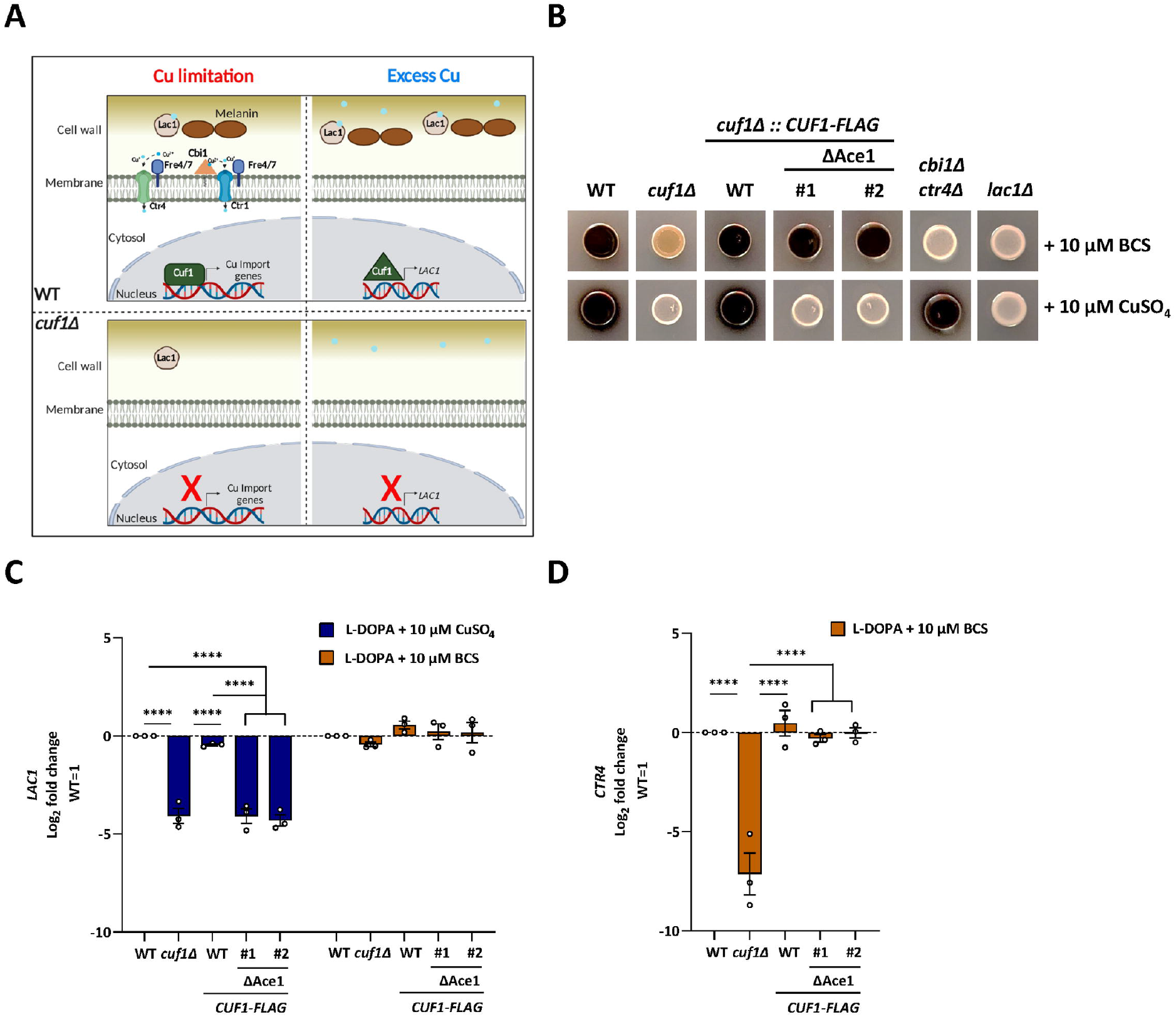
Cuf1 Ace1-like Cys-rich motif is essential for LAC1 regulation in response to excess copper. **(A)** Schematic summary of the role of Cu and Cuf1 regulation of Lac1 activity and melanin production during both Cu limitation and Cu excess. In the absence of Cuf1-induction of Cu-import genes during Cu limitation, melanin production by Lac1 is impaired. Cuf1 is also required for expression of the *LAC1* gene during Cu excess. Images made using BioRender. **(B)** Melanization during Cu limitation (10 μM BCS) and in presence of exogenous copper (10 μM CuSO_4_). Indicated strains were grown for 18 h in SC at 30°C, diluted to OD_600_ of 2.5 and spotted onto L-DOPA agar plates with indicated supplementation. L-DOPA plates were incubated for 3 days at 30°C. Melanin production is demonstrated by colony pigmentation **(C-D)** *LAC1* (C) and *CTR4* (D) transcript abundance in melanin-inducing conditions. Indicated strains were diluted to OD_600_ of 0.5 in liquid L-DOPA medium supplemented with either 10 μM BCS or 10 μM CuSO_4_ and grown for 3 h at 30°C. Relative transcript abundance for *LAC1* (C) or *CTR4* (D) was assessed by quantitative RT-PCR and normalized to WT transcript levels via the ΔΔC_T_ method. The data is presented as the mean +/-SEM of the log_2_ fold change of 3 biological replicates. Statistical analysis was performed using a one-way ANOVA.

The WT strain was strongly melanized by 72 h, and the addition of excess Cu to the medium resulted in a higher degree of pigmentation (Fig 4B). In contrast, the *cuf1*Δ mutant displayed decreased melanin in Cu-limited conditions, similar to that of the *cbi1*Δ *ctr4*Δ Cu import mutant (Fig 4B)[28, 46]. In these conditions, *LAC1* gene expression in the *cuf*1Δ mutant was similar to WT, suggesting that the observed decreased melanin formation in the *cuf1*Δ mutant is due to failed induction of Cu import genes, such as *CTR4*, resulting in reduced levels of intracellular Cu (Fig 4C-D). Consistent with this hypothesis, we observed full restoration of melanization in the Cu-import mutant with Cu supplementation, restoring Lac1 Cu loading and thus enzymatic function (Fig 4B).

In contrast to the Cu import mutant, melanin formation was not restored in the *cuf1*Δ mutant when the medium was supplemented with Cu. On the contrary, the *cuf1*Δ strain demonstrated an even more severe melanin-deficient phenotype, similar to that of the *lac1*Δ mutant (Fig 4B). In contrast to intact *LAC1* expression during Cu-limitation, we observed a marked relative decrease in *LAC1* transcript levels in the *cuf1*Δ mutant strain in Cu-excess conditions, consistent with the role of Cuf1 supporting *LAC1* expression when Cu is present in excess. (Fig 4C).

Reintroduction of the *CUF1-FLAG* WT allele fully restored WT levels of melanin production in both high and low Cu environments, as well as WT levels of *LAC1* and *CTR4* expression (Fig 4B-D). In contrast, the *cuf1*Δ strain expressing the ΔAce1-like Cys site mutant allele only restored melanization when Cu was limited but not when Cu was present in excess (Fig 4B). In this strain, *CTR4* expression was fully restored in Cu-limiting conditions, but *LAC1* expression was not restored in Cu excess (Fig 4C-D). Taken together, these data demonstrate a major role for Cuf1 in controlling melanin formation, by ensuring adequate levels of Cu as catalytical cofactor for Lac1 in low Cu conditions, and by supporting *LAC1* gene expression in higher Cu conditions. Additionally, the Cuf1 Ace1-like Cys-rich motif is required for ensuring proper *LAC1* expression and melanization when Cu is present in excess but is not required for melanization during Cu limitation.

#### The effect of *Cn*Cuf1 on superoxide dismutase (SOD) expression and ROS detoxification

The Cu, Zn-superoxide dismutase Sod1 helps to protect the fungal cell from ROS generated by host immune cells during infection [26]. Previous studies demonstrated that Cuf1 binds to the *SOD1* promoter and induces *SOD1* gene expression in high Cu stress. In contrast, during prolonged Cu starvation, *SOD1* expression is repressed to divert limited Cu from this Cu-containing enzyme to support other Cu-dependent cell processes. In this condition, Cuf1 binds to the *SOD2* promoter and induces the expression of a cytosolic retained variant of this Mn-dependent mitochondrial superoxide dismutase to uncouple ROS detoxification from a Cu-requiring enzyme when cellular Cu is scarce [11]. Therefore, deletion of *CUF1* renders cells vulnerable to ROS stress not only during high Cu stress due to failure of up-regulation of *SOD1* but also during low Cu stress due to inability to switch to a Cu-independent ROS detoxification pathway (Fig 5A).

**Figure 5:**
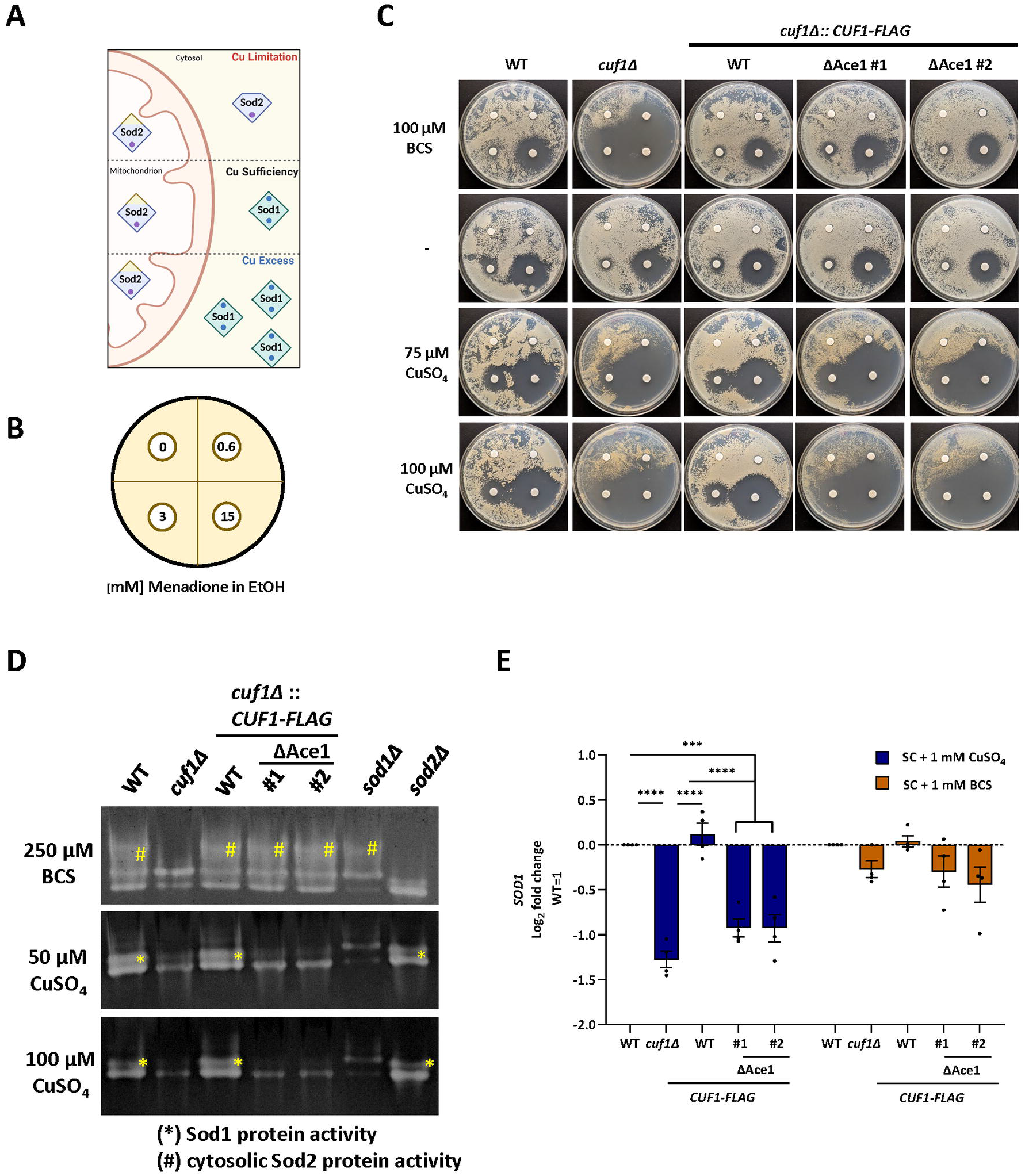
Cuf1 Ace1-like Cys-rich motif is required for superoxide dismutase (Sod)-dependent ROS detoxification during high Cu stress. **(A)** Schematic summary of the role of Cu and Cuf1 for Sod protein regulation during Cu limitation and excess. Cu, Zn-containing Sod1 is the major cytosolic Sod in Cu sufficiency and excess. The mitochondrial Mn-dependent Sod2 assumes the majority of cytosolic Sod function during Cu limitation. Images were made using BioRender. **(B-C)** Disc diffusion assay to test cellular superoxide sensitivity. Indicated strains were inoculated on SC agar plates supplemented with indicated amounts of BCS or CuSO_4_. Cells were challenged with increasing concentrations of the superoxide generator menadione on filter discs as depicted in (B) for 3 days at 30°C (C). **(D)** In-gel SOD activity assays from whole cell extracts of indicated strains. For Cu stress induction, strains were incubated in SC medium supplemented with indicated amounts of BCS or CuSO_4_ for 24 h at 30°C. Whole cell extracts were generated, separated by native PAGE, and stained for SOD activity using nitroblue tetrazolium (NBT) staining solution. **(E)** Transcript abundance of the *SOD1* gene during high Cu and low Cu stress. Indicated strains were diluted to OD_600_ 0.5 and incubated for 2 h at 30°C in SC medium. Cu stress was induced by the addition of either 1 mM BCS or 1 mM CuSO_4_ and cultures were incubated for 3 h at 30°C. Relative transcript abundance for *SOD1* was assessed by quantitative RT-PCR and normalized to WT transcript levels using the ΔΔC_T_ method. The data is presented as the mean +/-SEM of the log_2_ fold change of 3 biological replicates and statistical analysis was performed using a one-way ANOVA.

To test the effects of the Ace1-like Cys-rich motif mutation on cellular ROS detoxification, we incubated the WT and *cuf1*Δ strains, as well as strains expressing *CUF1-FLAG* WT or the ΔAce1-like Cys mutant allele, in the presence of increasing concentrations of the superoxide-generating compound menadione by disc diffusion (Fig 5B). Consistent with previous studies, the *cuf1*Δ mutant showed an increased susceptibility to menadione, measured by zone size of growth inhibition, in both high and low Cu conditions (Fig 5C) [11]. Both Cu-stress phenotypes were fully restored in the *cuf1*Δ strain expressing the *CUF1-FLAG* WT allele. In contrast, the strain expressing the *CUF1-FLAG* ΔAce1-like Cys mutant allele only restored ROS resistance during low Cu stress but not high Cu stress (Fig 5C).

To fully analyze the role of the Ace1-like Cys motif in regulating SOD-driven ROS detoxification, we measured Sod activity in whole cell extracts using an in-gel Sod activity assay [10] from cells conditioned in either low or high Cu stress. The Sod activity gels demonstrated three activity bands from cell extracts from the WT strain incubated in low Cu conditions. The lowest band likely represents Sod1, while the middle band likely represents Sod2 activity given their absence in the respective *sod1*Δ and *sod2*Δ mutants (Fig 5D). The highest of the three bands was absent in the *cuf1*Δ and *sod2*Δ mutant strains (Fig 5D, yellow hash tag), and likely represents the cytosolic Sod2 variant that is specifically enriched in Cu limitation in a Cuf1-dependent manner [11]. This Sod activity is restored in the *cuf1*Δ mutant by introduction of either the *CUF1-FLAG* WT or the ΔAce1-like Cys site mutant allele, indicating that Cuf1-dependent formation of cytosolic Sod2 during Cu-limiting conditions is not dependent on the Cuf1 Ace1-like Cys motif (Fig 5C-D).

The patterns of SOD activity varied when the strains were conditioned in high Cu stress, with two predominant bands of similar intensity in WT cells. Both bands were significantly reduced in intensity in the *cuf1*Δ mutant strain. Moreover, in the *sod1*Δ strain, these two bands were only faintly present, and the upper of these two was either not present or altered in its molecular size (Fig 5D, yellow asterix), suggesting that this upper band represents Sod1 activity. While expression of *CUF1-FLAG* WT restored the Sod activity bands to WT-like levels, we observed identical patterns of reduced Sod activity in the *cuf1*Δ mutant as well as the *cuf1*Δ mutant expressing the *CUF1-FLAG* ΔAce1-Cys mutant allele, indicating that the first Cys-rich motif is required for Cuf1-depenedent induction of Sod1 protein activity in response to high Cu stress (Fig 5D). Consistent with this finding, the Cuf1 Ace1-like Cys-rich motif is required for supporting Cuf1-dependent *SOD1* transcript induction in high Cu conditions (Fig 5E). Together, these phenotypic data, enzyme activity assays, and transcript assessments demonstrate the importance of the Ace1-like Cys-rich motif for Cuf1-dependent regulation of ROS detoxification in a high Cu stress environment.

### Consequences of a blunted Cuf1 high Cu stress response on fitness, yeast containment and immune response inside the host lung environment

A previous study conducted by Waterman *et al*. established a role for Cuf1 for virulence in an intravenous model of *Cn* infection, demonstrating a decreased fungal burden of the *cuf1*Δ mutant compared to WT in the mouse brain but not in the mouse lung [32]. More recent studies have suggested that these different anatomic sites represent different Cu environments encountered by *Cn* during the course of infection [21, 22, 28]. Using high and low Cu promoter-driven Luciferase strains, one study demonstrated an up-regulation of genes typically induced during Cu deficiency in the brain, while the lung environment induced genes required for Cu detoxification [21]. This finding was further underscored by assessment of the effects of Cu homeostasis mutant strains on virulence and survival at the different sites of infection: strains lacking components of the Cu import machinery were less fit within the mouse brain environment, while strains lacking components required for Cu detoxification were less fit in the mouse lung environment [21, 22, 28].

Given the high Cu stress phenotypes observed *in vitro* for the *cuf1*Δ mutant and the ΔAce1-like Cys-rich motif mutant strain, we hypothesized that these strains would display altered physiology within the high Cu environment of the lung. We inoculated female A/J mice by inhalation with the WT strain, the *cuf1*Δ mutant, or the *cuf1*Δ mutant reconstituted with either the WT *CUF1-FLAG* allele or the *CUF1-FLAG* ΔAce1 mutant allele. Lungs were harvested at 14 days post infection, a time point at which lung infection is well established but before animals succumb to infection. In the infected lungs, we assessed lung fungal burden as well as relative patterns of inflammation by histology and flow cytometry analysis (Fig 6, Supp. Fig 3). Consistent with prior studies, we did not observe significant difference in lung fungal burden for any of the analyzed strains at this infection time point (Fig 6A) [32]. However, we did note a significant increase in lung wet weight in animals infected with the WT or *CUF1-FLAG* WT strain compared to the lungs of mice infected with the two high Cu response-defective strains (Fig 6B). Additionally, we observed macroscopic differences between WT and *CUF1-FLAG* WT-infected lungs compared to *cuf1*Δ- and the *CUF1-FLAG* ΔAce1-like Cys-rich motif mutant-infected lungs: WT-infected lungs demonstrated more extensive foci of tissue abnormality (darkening) compared to lungs infected with either the *cuf1*Δ mutant or the ΔAce1-like Cys-rich motif mutant strain (Fig 6C, white arrows).

**Figure 6:**
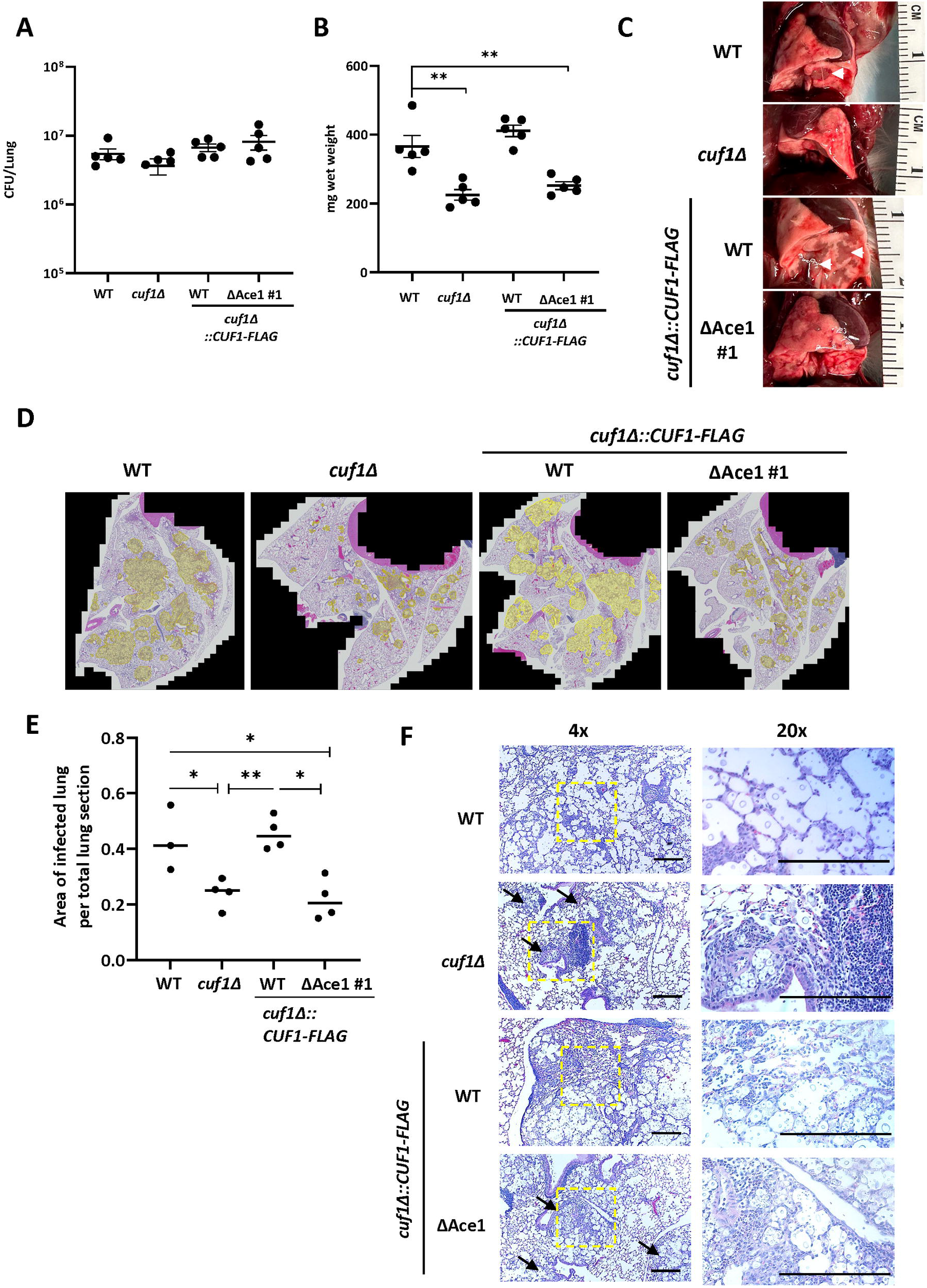
Effects of Cuf1 high Cu stress regulation on survival, yeast cell dissemination and inflammation in the infected mouse lung. Intranasal infection of female A/J mice. Mice were infected by inhalation with 10^5^ cells of the indicated strains (5 mice/strain). Mice were sacrificed 14 days postinfection and their lungs harvested. **(A-C)** Lung fungal burden 14 days postinfection. Lung lysates were assessed by quantitative culture (A), lung wet weight (B) and macroscopic lung abnormalities (C). (A-B) Data was plotted, and an unpaired t-test was performed using GraphPad Prism (B only). (C) Foci are indicated with white arrows. **(D-F)** Histopathology analysis of infected murine lungs 14 days post infection. Mice were sacrificed and lungs were fixed via intratracheal instillation of neutral buffered formalin under gravity flow. Hematoxylin and eosin staining were used to visualize microscopic lung pathology. (D) Representative stitched 20x tile scan images used for quantification of infected/inflamed lung area. Infected/inflamed areas are highlighted in yellow. (E) Quantification plot of infected versus non-infected section area using stitched 20x images. Plot was generated via GraphPad prism and a Welch’s t Test with unequal sample sizes was performed. (F) Representative 4x and 20x images. Yeast enriched foci of inflammation are highlighted by a black arrowhead. The length of the black bar represents 250 μm.

Histopathology analysis of fixed lung tissue demonstrated similarly altered patterns of inflammation and yeast containment between these two groups. The regions of lung tissues characterized by intense inflammation were demarcated and quantified relative to the total lung area. These measures demonstrated significantly more regionally extensive lung inflammation in mice infected with the WT strain or the *CUF1-FLAG* WT strain compared to either the *cuf1*Δ mutant or the Ace1-like *cuf1*Δ mutant strains (Fig 6D-E, yellow highlighted areas). While lungs infected with the WT strain displayed expected patterns of diffuse inflammation surrounding single and grouped encapsulated yeast cells throughout the lung tissue (Fig 6F) [47], the *cuf1*Δ mutant strain displayed yeast cells mostly confined to well circumscribed foci, similar to the early granulomas described previously in studies of *Cn* strains with slow growth phenotypes due to defects in mitochondrial function [47]. Also compared to WT, the *cuf1*Δ mutant-infected lungs displayed a very sharp distinction between inflamed regions containing encapsulated yeast cells and normal-appearing lung tissue with no apparent yeast cells present (Fig 6F).

Lungs infected with the *CUF1-FLAG* WT-expressing strain demonstrated similar patterns of inflammation as observed in WT-infected lungs (Fig 6D-F). In contrast, the *CUF1-FLAG* ΔAce1-like Cys-rich motif mutant expressing strain generated an infection pattern similar to *cuf1*Δ-infected lungs, with smaller and more contained areas of inflammatory cells intermixed with yeast cells surrounded by normal-appearing lung tissue with no apparent yeast cells (Fig 6D-F). These findings suggest that Cuf1-regulated high Cu responses are not required for absolute fitness within the mouse lung environment but instead modulate colonization patterns, yeast containment, and inflammation at the site of infection.

We also performed flow cytometry to test whether the different lung infection patterns observed histologically in WT-versus *cuf1*Δ mutant-infected lungs were marked by changes in the immune cell populations. Despite the observed macroscopic and microscopic differences between the WT- and mutant-infected lungs, we noted no significant differences at this 14-day infection time point in the relative percentages of immune cell subtypes (Total Myeloid cells - Neutrophils, Monocytes, Eosinophils, and Dendritic cells as well as Total Lymphocytes - B cells and T cells) or T cell activation markers (IFN γ, IL-17A and IL-4 measured in CD4^+^, CD8^+^ and γΔ T cells, Supp. Fig 3).

## Discussion

*Cn*Cuf1 is unique among fungal Cu-responsive transcription factors as it regulates a diverse spectrum of Cu responses, combining the role of a high Cu-responsive transcription factor with the role of a low Cu-responsive transcription factor [23]. Accordingly, it shares conserved motifs with both high and low Cu-responsive transcription factors previously identified in model ascomycete yeast Cu transcription factors [23]. Previous research in these ascomycete model yeasts has demonstrated that Cu binds directly to these Cys-rich motifs and thus modulates protein confirmation. These confirmational changes result in either activation (Ace1-like high Cu-responsive TFs) or inhibition of Cu-responsive transcriptional activity (Mac1-like low Cu-responsive TFs) [35, 36, 38, 39, 48].

While these regulatory Cys-rich motifs are conserved in fungal Cu-responsive TFs, their precise molecular function for regulation of Cu-responsive transcriptional activity has evolved differently in different fungal clades. For instance, an interspecies cross-complementation study conducted with the *Aspergillus fumigatus* (*Af*) AceA high Cu-responsive transcription factor demonstrated that AceA homologs from ascomycete yeasts did not functionally complement the high Cu growth defect observed in the *Af* Δ*aceA* mutant, suggesting that, even though the N-terminal DNA binding and the Cu sensing motifs are highly conserved among these high Cu-responsive transcription factors, the motifs might function differently in *Aspergillus* species than in their yeast counterparts [49]. Another example for this divergence is the Cu-responsive TF, *Sp*Cuf1, found in fission yeast, which encodes for an Ace1-like Cys-rich motif and an additional Mac1-like Cys-rich motif. Here, Cu binding to its Cys-rich motifs does not only result in transcriptional inhibition but also cytosolic re-localization of the TF when Cu is in excess [33, 34, 50]. Among the characterized fungal Cu-responsive TFs *Cn*Cuf1 demonstrates the largest divergence, having evolved into a dual Cu-stress responsive transcription factor, which directly binds to and regulates different subsets of target genes in response to low and high Cu stress. *Cn*Cuf1 is not only the first described dual Cu-responsive TF, but it also regulates Cu stress adaptation of a basidiomycete fungus, which is distantly related to the well-studied model ascomycetes. Our research reinforces this evolutionary divergence; while the Ace1-like conserved Cys-rich motif maintained its functional relevance for *Cn*Cuf1 high Cu stress adaptation, the Mac1-like Cys-rich motifs lost their predominant role in regulating low Cu stress adaptation. Furthermore, the absolute dependence on the presence of the unstructured *Cn*Cuf1 C-terminal domain for Cu stress sensing and adaptation further underscores the evolutionary reshaping of molecular mechanisms driving *Cn*Cuf1 transcriptional activity. Additional research will be necessary to fully understand the molecular mechanisms behind the *Cn*Cuf1 Cu-dependent transcriptional activity, and further phylogenetic studies will be required to define the diversity of Cu stress sensing within the fungal kingdom.

The transition metal Cu has been established as an important virulence factor in all major human fungal pathogens as it is used as a cofactor for several enzymes required for stress adaptation within the host [10, 11, 42, 51, 52]. Accordingly, Cu import genes and the Cu-responsive transcription factors sensing and maintaining the cellular Cu quota have been demonstrated to play an important role in fungal pathogenesis [12, 22, 28, 32, 53-55]. For instance, in *Candida albicans* (*Ca*) researchers demonstrated that the low Cu-responsive *Ca*Mac1 transcription factor is required for host invasion, adhesion to host cells, and antifungal tolerance [53]. Similarly, mutation of the low Cu-responsive MacA transcription factor in *Af* attenuates pathogenicity in a neutropenic immunocompromised mice model [54].

Nutritional immunity responses for other transition metals mostly involve sequestering these micronutrients from the sites of infection [15, 17]. In contrast, Cu nutritional immunity responses appear to be more dynamically regulated and are highly tissue dependent. While Cu might be restricted at one site or time point of an infection, elevated and toxic Cu levels are present at other sites or time points to act as an antimicrobial agent to kill the invading pathogen [10-13, 16, 21, 22, 56]. Therefore, high Cu stress responses are as important as Cu scavenging responses for microbial pathogenesis. However, both responses must be balanced to ensure proper adaptation to the Cu environment established at the particular infection micro-niche.

Analysis of the dynamic shifts in the host and microbial physiology associated with Cu nutritional immunity responses has been challenging to study in *Cn* since deletion of the *CUF1* gene ultimately blunts both high and low Cu stress responses [23]. This makes deciphering the role of high and low Cu stress responses during infection difficult as these stresses can have compensatory effects. For instance, deletion of the *CTR4* Cu importer gene was shown to be beneficial for *Cn* survival during pulmonary infection, preventing an excessive influx of Cu and thus reducing the toxic effect of the high Cu lung environment. In contrast, the same gene deletion resulted in reduced fungal virulence when the strain was directly injected into the CNS, a site where Cu is limited [22]. Our mutational study aimed to identify amino acid motifs specifically required for transcriptional activity in high or low Cu environments to uncoupled *Cn*Cuf1-driven high Cu stress responses from low Cu stress responses. In this work using a combination of transcriptional analysis, Cu stress phenotypic analysis, and enzyme activity assays, we established an essential role for the Ace1-like Cys-rich motif for *Cn*Cuf1 transcriptional activity during high Cu stress. Importantly, mutation of this site does not affect low Cu stress adaptation. Using this site mutant, we were able to demonstrate the consequences of a blunted high Cu stress adaptation on Cu-dependent virulence factor expression *in vitro* and on fungal growth, dissemination and inflammation *in vivo*.

Recent work conducted with *Af*AceA demonstrated that mutation of its conserved Cys-rich region attenuates virulence of the fungus similar to a complete Δ*aceA* mutation [49], highlighting the importance of this Cys-rich motif for activating Cu stress responses within the mouse lung environment. Similar to a Δ*aceA* mutant in *Af*, deletion of the *CnCUF1* gene results in an attenuation of virulence [32]. However, unlike the *Af* Δ*aceA*, a *CUF1* deletion did not affect *Cn* growth inside the lung environment [32, 49]. This observation stands in contrast with the severe *in vitro* phenotypes exhibited by the *cuf1*Δ mutant and the *CUF1-FLAG* ΔAce1 motif mutant strains during high Cu stress. The observed *in vivo* growth phenotype of the *cuf1*Δ and the *CUF1-FLAG* ΔAce1 mutant is also very different from the strongly attenuated phenotype observed for a *Af* Δ*aceA* mutant. These very distinct outcomes of a blunted high Cu stress response observed in the pathogenesis of two fungal pathogens in the same host organ suggest different host-pathogen dynamics within the lung Cu environment, which are likely dependent on the type of pathogen and its lifestyle inside the host tissue. For example, the hyphal fungus *Af* exists primarily extracellularly in the infected lungs, whereas the yeast-like *Cn* survives both intracellularly within host immune cells and extracellularly in the lung tissue. Another possibility is that *Cn* might not be as dependent on Cuf1 as *Af* on AceA for Cu detoxification *in vivo*. For instance, *Cn* could have evolved alternate compensatory mechanisms to survive in the host lung environment which are independent from Cuf1. To address these questions and to better understand the dynamics of Cu nutritional immune responses within the lung environment, a more thorough assessment is needed at several different stages of infection.

Although we did not observe significant differences in fungal burden between our WT and mutant strains at two weeks of infection, we did observe macroscopic and microscopic differences within the lungs of WT-infected mice compared to mice infected with the *cuf1*Δ mutant or *CUF1-FLAG* ΔAce1 site mutant. This observation suggests a role for *Cn*Cuf1-driven high Cu stress responses for modulating inflammatory responses within the lung environment, by potentially mediating adaptive responses at the fungal cell surface. This fungal feature is composed of ordered layers of polysaccharides and mannosylated proteins and is the site of recognition and interaction with the host immune system. Changes in carbohydrate composition, as well as masking or unmasking of certain cell surface components, can modulate and define the pattern of inflammation caused by the pathogen [57-59]. For instance, chitin has been associated with non-protective immune responses, and the degree of chitin masking, coupled with the relative amount of chitosan within the *Cn* cell wall, modulates the severity of immune recognition and inflammation within the host tissue [59, 60].

Previous work has suggested that the fungal cell wall also plays a role in regulating intracellular Cu levels during Cu stress [27]. In turn, *Cn*Cuf1 regulates multiple genes associated with the fungal cell wall in response to Cu stress [23, 27]. Another study showed that cell wall architecture and cell wall chitosan levels are significantly altered in a *Cn* Cu homeostasis mutant strain during Cu limitation [27]. These findings document a role for Cu and *Cn*Cuf1 in modulating the fungal cell wall. Hence, one possible explanation for the observed altered macroscopic and microscopic phenotypes could be that the impaired high Cu stress response in these mutant strains prevented Cu-stress-driven cell wall remodeling and thus modulated immune recognition and yeast dissemination, which could trigger a more focal, granuloma-like inflammatory response with fungal containment within these inflammatory foci. However, we did not observe any significant changes of the relative percentages of immune cell subtypes or T-cell activation, which could explain the altered microscopic features found in mutant-infected lungs.

Another possible explanation of the altered yeast distribution observed in mutant-infected lungs could be the presence of different Cu nutritional micro-niches restricting dissemination of yeast cells with an impaired high Cu stress adaptation to lung micro-niches presenting a more favorable, non-toxic Cu environment. This model is supported by previous research demonstrating the mouse lung to be a high Cu environment, which induces *Cn* Cu detoxification genes [21, 22]. However, no clear data is available which lung micro-niches might induce the *Cn* high Cu stress responses. Further investigations will more precisely define the interplay between Cu availability within different host micro-niches, the effects of Cu and *Cn*Cuf1 on the fungal cell surface and their impact on fungal dissemination and immune recognition at different stages and sites of infection.

## Materials and Methods

### Ethics statement

All animal experiments conducted in this study were approved by the Duke University Institutional Animal Care and Use Committee (IACUC) (protocol #A102-20-05). Mice were handled according to IACUC guidelines.

### Strains, media and growth conditions

*Cryptococcus neoformans* strains used in this study are shown in **Table S1**. Strains generated in this study were created using the *C. neoformans var. grubii H99* background. Strains were created using biolistic transformation or CRISPR/Cas9 method as previously described [61, 62]. Used guide RNA for CRISPR/Cas-based transformations are listed in **Table S2**. Yeast extract (2%)-peptone (1%)-dextrose (2%) (YPD) medium supplemented with 2% agar, 200 μg/ml of hygromycin B (HYG) and 10 µM CuSO_4_ was used for colony selection after transformation. Transformants were screened by PCR for intended mutations and gene expression was confirmed by western blot. Plasmids and oligonucleotides used in this study for strain generation are listed in **Tables S3-S4**. For strains generated through mating, the corresponding *MAT* a and *MAT* α strain were mixed on to a MS agar and incubated in the dark until basidiospores were formed. Dissected spores were grown on YPD media at 30C for 3-4 days and screened for marker, gene expression (via western blot) and parental Cu-stress phenotypes.

Strains were stored and recovered from glycerol stocks (stored at ™80°C) and were routinely cultivated in synthetic complete (SC) medium (MP Biomedicals) at 30°C. Media was supplemented with indicated concentrations of CuSO_4_ or the extracellular Cu^+^ chelator bathocuproine disulfonate (BCS) to induce Cu excess, sufficiency or deficiency, respectively.

Growth phenotype analysis was performed as previously described [27, 63]. Melanization assays were performed on L-3,4-dihydroxyphenylalanine (L-DOPA) media (7.6 mM L-asparagine monohydrate, 5.6 mM glucose, 22 mM KH_2_PO_4_, 1 mM MgSO_4_ heptahydrate, 0.5 mM L-DOPA, 0.3 μM thiamine-HCl, 20 nM biotin, pH 5.6) as previously described [27, 63]. L-DOPA plates were incubated at 30°C for 3 days.

### Trichloroacetic acid (TCA)-based protein precipitation and western blotting

*Cn* cultures were grown overnight for 18 h in synthetic complete (SC) medium (MP Biomedicals) and were Cu-stressed as indicated. TCA-precipitated protein extracts were generated as previously described [64]. Precipitated proteins were solubilized in alkylating buffer (0.1 M Tris pH 8.3, 1 mM EDTA, 1% SDS) for 10 min at 65°C. For Cuf1-FLAG western blots, crude TCA extracts were used in all experiments and indicated amounts of the crude TCA extract were treated with 1x SDS-loading buffer (Biorad) and boiled for 10 min at 95°C. For Cu sensor blots, the crude TCA extract was clarified for 10 min at 21000 xg and total protein was quantified via BCA assay (Pierce). 20 µg total protein per sample was treated with 1x SDS-loading buffer (Biorad) and boiled for 10 min at 95°C. Protein samples were run on a 7.5% TGX Midi (Cuf1-FLAG) or 4-20 % Midi (Cu Sensor) polyacrylamide gel (Biorad) using SDS running buffer (Biorad) for ∼60 min at 150 V. Then, proteins were blotted on nitrocellulose using the Transblot Turbo system (Biorad) at the high molecular weight (Cuf1-FLAG site mutants), mixed molecular weight (Cuf1-FLAG fragment) or standard (Cu sensor) midi blotting setting. Protein transfer was validated via ponceau staining and the membrane was blocked for 1 h at room temperature (RT, for primary antibodies) or overnight at 4°C (for HRP-conjugated antibodies) with 5% Milk in TBS + 0.1% Tween20 (TBS-T). Cuf1-FLAG protein levels were analyzed using a monoclonal Anti-FLAG M2-Peroxidase antibody (1:5000 dilution) (Sigma, α-FLAG-HRP) using a rabbit anti-GAPDH at 1:2000 dilution in 5% milk + TBS-T and 0.02% sodium azide (Abcam, ab9485) combined with an anti-rabbit-HRP in a dilution of 1:4000 (GE Healthcare) as loading control. For mClover detection, a mouse anti-GFP at 1:5000 dilution in 5% milk + TBS-T and 0.02% sodium azide (Roche, 11814460001) combined with an anti-mouse-HRP in a dilution of 1:4000 (Amersham, NA931V) was used. For mRuby detection, a rabbit anti-RFP at 1:1000 dilution in 5% milk + TBS-T and 0.02% sodium azide (Abcam, ab124754) combined with an anti-rabbit-HRP in a dilution of 1:4000 (Amersham, NA934V) was used. As loading control, a rabbit anti-GAPDH antibody was used as described above. Primary antibodies were incubated overnight at 4°C, the FLAG-HRP conjugate as well as secondary antibodies were incubated for 1 h at RT. Blots were blocked with 5% milk in TBS-T and washed 3x for 15 min with TBS-T after each antibody incubation.

For detections using multiple primary antibodies, the membrane was stripped using the Restore Western Blot Stripping Buffer (Thermo Fisher Scientific) according to manufacturer’s instructions and re-blocked for 30 min in between each primary antibody detection. HRP-conjugated antibodies were detected by chemiluminescence using the DuraWest Pico substrate (Thermo Fisher Scientific). Chemiluminescence was measured using an Azure 600 imager (Azure Biosystems).

### RNA isolation and quantitative reverse transcriptase (qRT)-PCR

For transcript analysis during high versus low Cu stress, overnight cultures (∼18 h) incubated in synthetic complete (SC) medium (MP Biomedicals) were diluted to OD_600_ 0.5, grown for 2 h at 30°C in SC medium. Cu stress was then induced by the addition of either 1 mM BCS or 1 mM CuSO_4_ and cultures were grown for 3 h at 30°C.

For transcript analysis in melanin inducing conditions, overnight cultures grown in SC medium were diluted to OD_600_ 0.5 and incubated for 3 h at 30°C in L-DOPA medium supplemented with either 10 μM BCS or 10 μM CuSO_4_.

RNA extraction and cDNA analysis were performed as described previously [27]. For transcript analysis via qRT-PCR, cDNA was diluted 1:5 in RNase-free water and mixed with the PowerUP SYBR Green Master mix (Applied Biosystems) per protocol instruction and analyzed on a QuantStudio 6 Flex (Applied Biosystems). The oligonucleotides used for transcript analysis are listed in **Table S3**. Cycle threshold (C_T_) values were calculated using the QuantStudio 6 Flex and gene expression values were normalized to *GAPDH*. Expression fold changes were calculated using the ΔΔC_T_ method. For all transcript analysis a minimum of 4 independent biological replicates were analyzed.

### Chromatin immunoprecipitation followed by quantitative PCR (ChIP-PCR) analysis

Overnight cultures (∼18 h) grown in SC medium were diluted to OD_600_ 0.2, and incubated for 2 h at 30°C in SC medium. Cu stress was then induced by the addition of either 1 mM BCS or 1mM CuSO_4_ and cultures were grown for 2.5 h at 30°C. Crosslinking of the samples was performed in a 1% formaldehyde solution for 30 min at RT. The downstream processing and ChIP were performed as previously described [23] with the following adaptions: Instead of sonication 5 times, with 30 sec ON followed by 1 min OFF, samples were processed using a Bioruptor UCD200 (Diagenode) via 24 cycles of 15 sec “ON “ followed by 15 sec “OFF” at the high level to shear chromatin. Immune precipitated DNA fragments were purified with a PCR clean up kit (Macherey Nagel) following manufacturer’s instructions. qPCR was performed with the PowerUP SYBR Green Master mix (Applied Biosystems) per protocol instructions and analyzed on a QuantStudio 6 Flex (Applied Biosystems) using oligonucleotides listed in **Table S3**. For quantification, the C_T_ from each input was subtracted from the C_T_ from the target gene obtained for the Cuf1–FLAG strain (ΔC_T target_) or the negative control (WT strain, no FLAG-tagged Cuf1, ΔC_T control_). Then, the Δ C_T control_ was subtracted from the ΔC_T target_ to give a ΔΔ C_T_ value. For each gene promoter, Cuf1 enrichment was calculated as the inverse log_2_ of ΔΔ C_T_ (2^-^ΔΔ^Ct^). Cuf1-FLAG enrichment at the tubulin promoter was used for normalization.

### Disc diffusion assay

To measure menadione-induced ROS stress sensitivity, SC overnight cultures (∼18 h) of indicated *Cn* strains were diluted 1:500 in SC medium, and 500 μL cell suspension spread onto SC plates or SC plates supplemented with indicated amounts of BCS or CuSO_4_. Inoculated plates were dried for 45 min at RT. Filter discs were spotted with 15 μL of the indicated concentrations of menadione (dissolved in ethanol) and set in quadrants around the plates. Plates were incubated at 30°C for 3 to 4 days.

### Protein extraction and Sod activity assay

For Sod activity analysis, *Cn* overnight cultures (∼18 h) incubated in SC medium were diluted to OD_600_ 0.1 in SC medium or SC medium supplemented with either 250 μM BCS or 50 μM or 100 μM CuSO_4_. Cultures were incubated for 24 h at 30°C. For protein extraction, cells were pelleted (1 min, 11000 xg, 4°C), washed 1x with cold Lysis buffer (10 mM sodium phosphate, 50 mM NaCl, 0.1% triton X-100, 10% glycerol pH 7.8 made in chelex-treated H_2_O) and resuspended in 350 μL of cold lysis buffer + 1x Protease Inhibitor cocktail (ROCHE) and PMSF. Cells were lysed with a Bead Ruptor 12 (Omni international) using 6 cycles of 60 sec “ON” and 90 sec “OFF”, 5.5 speed at 4°C. Cell lysates were clarified for 15 min at 21000 xg at 4°C and the clarified lysate was transferred into a 1.5 mL reaction tube. Total protein levels were measured using a BCA kit (Thermo Fisher Scientific) according to manufacturer’s instruction. Lysates were diluted to a concentration of 75 μg protein in 25 μL native sample buffer (Bio-Rad, catalog #1610738) and lysis buffer added to a final volume of 50 μL. Diluted lysates at a volume of 20 μL were loaded onto a 10% mini-PROTEAN TGX precast protein gel (Bio-Rad, catalog #4561033). Electrophoresis was performed in 1x tris glycine running buffer at 200 V and 4°C for approximately 1 h. Gels were then incubated at room temperature (RT) in the dark for 1 h in 50 mL NBT staining solution (50 mM potassium phosphate, 30 μM riboflavin, 0.48 mM nitro blue tetrazolium chloride, pH 7.8) while shaking at 60 rpm. NBT staining solution was decanted and gels were soaked in diH_2_O in the light for approximately 30 min before imaging on an iBright 1500 FL Imager (Thermo Fisher Scientific).

### Infection experiments

We used the murine inhalation model of cryptococcosis [64] to assess fungal survival and host response after lung infection. For each strain, 5 female A/J mice were used. Mice were anesthetized by isoflurane inhalation and intranasally inoculated with 1 × 10^5^ fungal cells of PBS-washed overnight cultures (∼18 h) of indicated strains grown in SC. Mice were sacrificed at 14 days post infection and the mouse lung harvested, weighed, and homogenized in cold PBS. Colony forming units (CFU) were calculated by quantitative culture and are represented as CFU/lung. For Histopathology at 14 days post infection, mice were sacrificed using a lethal dose of pentobarbital sodium, and lungs were fixed via intratracheal instillation of neutral buffered formalin under gravity flow [65]. Fixed lungs were submitted to the Immunohistopathology Core at Duke University for paraffin embedding and staining with hematoxylin and eosin.

### Quantification of infection area in mouse lung sections

Whole images of the hematoxylin and eosin-stained mouse lung sections were obtained on a Keyence BZ-1000 in a 20x tile scan and stitched using Keyence Analysis software. Images were exported as OME TIFFs and passed through ‘bfconvert’ from the Bio-Formats command line tool (v. 8.3.0) to create image resolution pyramids. Images were then imported into QuPath (v. 0.6.0) for the remainder of analysis. Image scales were corroborated and 18 training regions were taken from the sample set of 15 slides to train a pixel classifier. Classes were used to identify ‘lung’ area vs. ‘non-lung’ and other structures. The entirety of the image was passed to the pixel classifier to identify the entirety of the lung area. This area was trimmed for mis-identification regions then passed to the classifier to add area measurements of defined classes, obtaining an area (m^2^) of lung parenchymal tissue excluding regions of blood and glass. In a blinded manner, samples were annotated for areas of infection based on the presence of (1) interstitial thickening, infiltration and distortion, (2) lymphocyte aggregates, and (3) bronchial/vessel cuffing with infiltrating cells. These annotation regions were then passed to the pixel classifier to calculate the area (m^2^) of lung parenchymal tissue within the infected regions. The infected/inflamed area divided by the total area of lung parenchyma yields the ratio of infection restriction.

### Processing lung tissue for single cell suspension for flow cytometry analysis

To prepare lung samples, mice were infected with yeast cells as described above and sacrificed at 14 days post infection. Lung cell harvest was performed as described previously [66]. Briefly, lungs were harvested, weighted and placed in 5 mL of lung digestion buffer (1x HBSS, 5% HI-FBS, 10 mM HEPES pH 7.2-7.4, 60 ug/mL Liberase (Roche, Cat# 5401119001), 6.5× 10^3^U/mL DNAseI (Millipore, Cat# 260913-10MU) and incubated for 25 min at 37°C and 225 rpm. Then, cells were passed through a pre-wetted (1x PBS+0.5% BSA) 70 µm mesh filter. Filtered cells were spun for 5 min (500 xg, 4°C) and supernatant was removed by decanting. The cell pellet was resuspended in 2 mL of hemolysis buffer (0.15 M ammonium chloride, 10 mM potassium bicarbonate, 1 mM EDTA, pH 7.2-7.4) and incubated at RT for 5 min. After incubation, the cell solution was neutralized by the addition of 10 mL 1x PBS to stop hemolysis and cells were pelleted (5 min, 500 xg, 4°C). Cells were resuspended in 1 mL 1x PBS+ 0.5% BSA and filtered through a pre-wetted 70 µm mesh filter as described above. The mesh filter was washed 2x with 1 mL 1x PBS + 0.5% BSA for a total volume of 3 mL per sample . Freshly isolated cells were used for all flow cytometry analysis, unless otherwise noted.

### Antibody staining for flow cytometry

For antibody staining, optimal cell density was determined by counting cells using a hemocytometer. Roughly 10^6^ cells per sample were analyzed. Cells were blocked using a 1:200 dilution of FC Block antibody (CD16/32, BioLegend Cat#101302) for at least 15 min before antibody staining. Antibodies used for flow cytometry analysis are listed in **Table S5**. Cells were incubated with antibodies for at least 15 min in the dark at RT, then washed with 1x PBS + 0.5% BSA (5min, 500 xg, 4°C). Cells were resuspended in resuspension buffer (for 1 sample: 5 uL Precision Counting Beads (BioLegend, Cat#424902), 1 uL DNAseI (Millipore, Cat# 260913-10MU), 94 uL 1x PBS + 0.5% BSA).

### Intracellular cytokine staining for flow cytometry

Single cell suspension were prepared from infected lungs as described above. To enhance cytokine production and detection, cells were plated to 2-3 × 10^6^ cells in 300 µL of cRPMI and 200 µL of 2.5x Phorbol-12-myristate-13-acetate (PMA)/ionomycin in cRPMI was added to cells (final concentrations per well: PMA (Millipore Sigma) 0.1 µg/mL; Ionomycin calcium salt(Adipogen) 0.375 µg/mL). Cells were incubated with the PMA/ionomycin mix for 30 min at 37°C and 5% CO_2_. Then, Brefeldin A (BioLegend-1x) was added to the cells and cells were incubated for up to 4 h at 37°C and 5% CO_2_. Following the 4 h incubation, cells were spun down (500 xg, 4°C, 5 min) and 250 µL of cell supernatant was removed. Then, cells were resuspended in 200 µL of residual media and transferred to a fresh staining plate. To assess cytokine production, cells were first stained with a mix containing Fc block (CD16/32, BioLegend, 1:200 dilution) and fixable LIVE/DEAD Zombie (ThermoFisher Scientific, 1:2000 dilution) for 10 min at 4°C in the dark. Extracellular staining followed for 30 min at 4°C in the dark. Next, cells were permeabilized and fixed with the Cyto-Fast Fix Perm Buffer Set (BioLegend) for intracellular cytokine staining following the manufacturer’s protocol. Briefly, cells were fixed by addition of 100 µL Cyto-Fast Fix Perm Buffer Set (BioLegend) at RT for 20 min. After fixation, cells were spun down (500 xg, 4°C, 5 min) and washed with perm/wash buffer (Cyto-Fast Fix Perm Buffer Set (BioLegend); 100 µL/well). Intracellular antibody staining was performed by adding 50 µL of antibody dilution in perm/wash buffer to the cells. Antibodies used for intracellular staining are listed in **Table S5**. The intracellular staining was performed in the dark at 4°C over night. Next day, cells were spun down (500 xg, 4°C, 5 min) to remove staining solution and washed with 100 µL of perm/wash buffer per well. The washed cells were resuspended in 100 µl 1x PBS.

### Flow cytometry analysis

The following markers were used to identify cell populations in the lungs: myeloid cells (CD45^+^, CD11b^+^), neutrophils (CD45^+^, CD11b^+^, Ly6G^+^), monocytes (CD45^+^, CD11b^+^, Ly6G^-^, Ly6C^+^), macrophages (CD45^+^, CD11b^+^, Ly6G^-^, CD64^+^, CD24^-^), exudate macrophages (CD45^+^, CD11b^+^, Ly6G^-^, CD64+, CD24^-^, Ly6C^+^,Siglec-F^-^), interstitial macrophages (CD45^+^, CD11b^+^, Ly6G^-^, CD64^+^, CD24^-^, Ly6C^-^, Siglec-F^-^), alveolar macrophages (CD45^+^, CD11b^+^, Ly6G^-^, CD64^+^, CD24^-^, Ly6C^-^, SiglecF^+^), eosinophils (CD45^+^, CD11b^+^, Ly6G^-^, CD24^+^, IA-IE^-^, SiglecF^+^), dendritic cells (CD45^+^, CD11b^+^, Ly6G^-^, CD24^+^, IA-IE^+^, DC), lymphocytes (CD45^+^,CD11b^-^), B cells (CD45^+^, CD19^+^, TCRβ^-^, CD11b^-^), T cells (CD45^+^, TCRβ^+^, CD3ε ^+^, CD19^-^, CD11b^-^), CD4^+^ T cells (CD45^+^, TCRβ^+^, CD3ε**+**, CD4^+^, CD19^-^, CD11b^-^), CD8^+^ T cells (CD45^+^, TCRβ^+^, CD3ε^+^, CD8^+^, CD19^-^, CD11b^-^), γδ T cells (CD45^+^, TCRβ^-^, CD19^-^, CD11b^-^, CD3ε ^+^, γδTCR^+^). Intracellular cytokines markers: IFNγ, IL-17A, and IL-4. Data was acquired on the Cytek Aurora (Cytek Biosciences) and data analyses was performed using Flow Jo (TreeStar).

## Statistical analysis

All data error bars represent standard errors of the means (SEM) of results from a number of biological replicates (N), as indicated in figure legends. Prior to statistical analysis, data from all experiments was log_2_ transformed for comparison of proportions. Statistical analysis was conducted using GraphPad Prism software v9. The statistical tests used in each experiment and their results (i.e., p values) are indicated in figure legends. Asterisks in figures correspond to statistical significance as follows: ****, P < 0.0001; ***, P = 0.0001 to P < 0.001; **, P = 0.001 to P < 0.01; *, P = 0.01 to P < 0.05;ns (not significant), P > 0.05.

## Acknowledgements

We thank the Duke Immunohistopathology Core for preparation and staining of the infected lung tissue. Research reported in this publication was supported by the National Institute of General Medical Sciences of the National Institutes of Health under award number R01GM145035 (C.D.J. and K.J.F.) and R01GM041840 (J.A.A. and C.P.) and by the National Institute Of Allergy And Infectious Diseases of the National Institutes of Health under award number R01AI184401 (J.A.A., C.P. and M.L.S) The content is solely the responsibility of the authors and does not necessarily represent the official views of the National Institutes of Health. J.A.A. and C.P. acknowledge further support by Duke University School of Medicine Bridge Funding 4532750. C.A.D.J received fellowship support from the Tri-Institutional Molecular Mycology and Pathogenesis Training Program (T32-Al052080).

