## Supplementary figures and images for "A cysteine-rich domain of the *Cryptococcus neoformans* Cuf1 transcription factor is required for high copper stress sensing and fungal virulence"

### Supp Fig 1

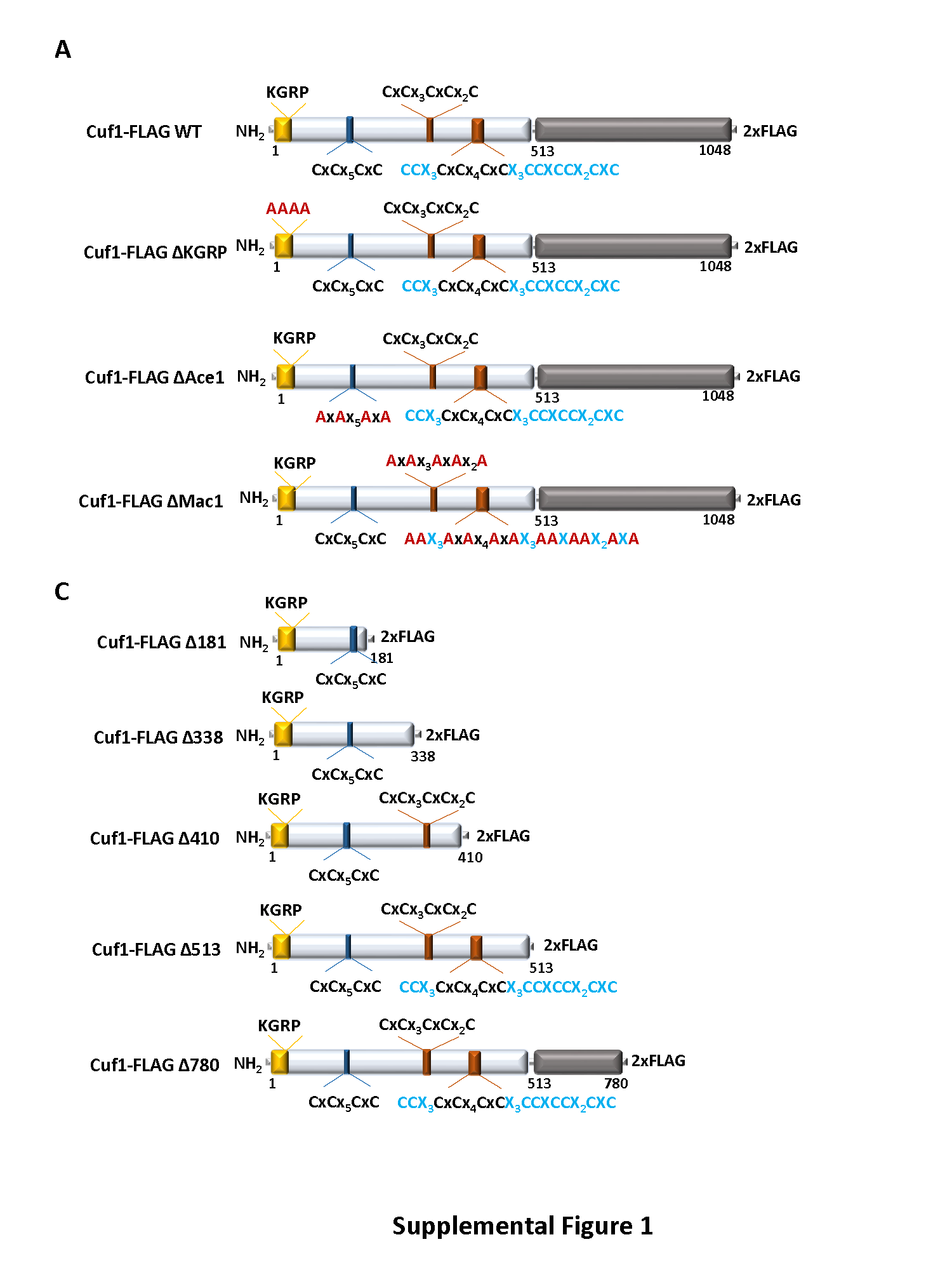

### Supp Fig 2

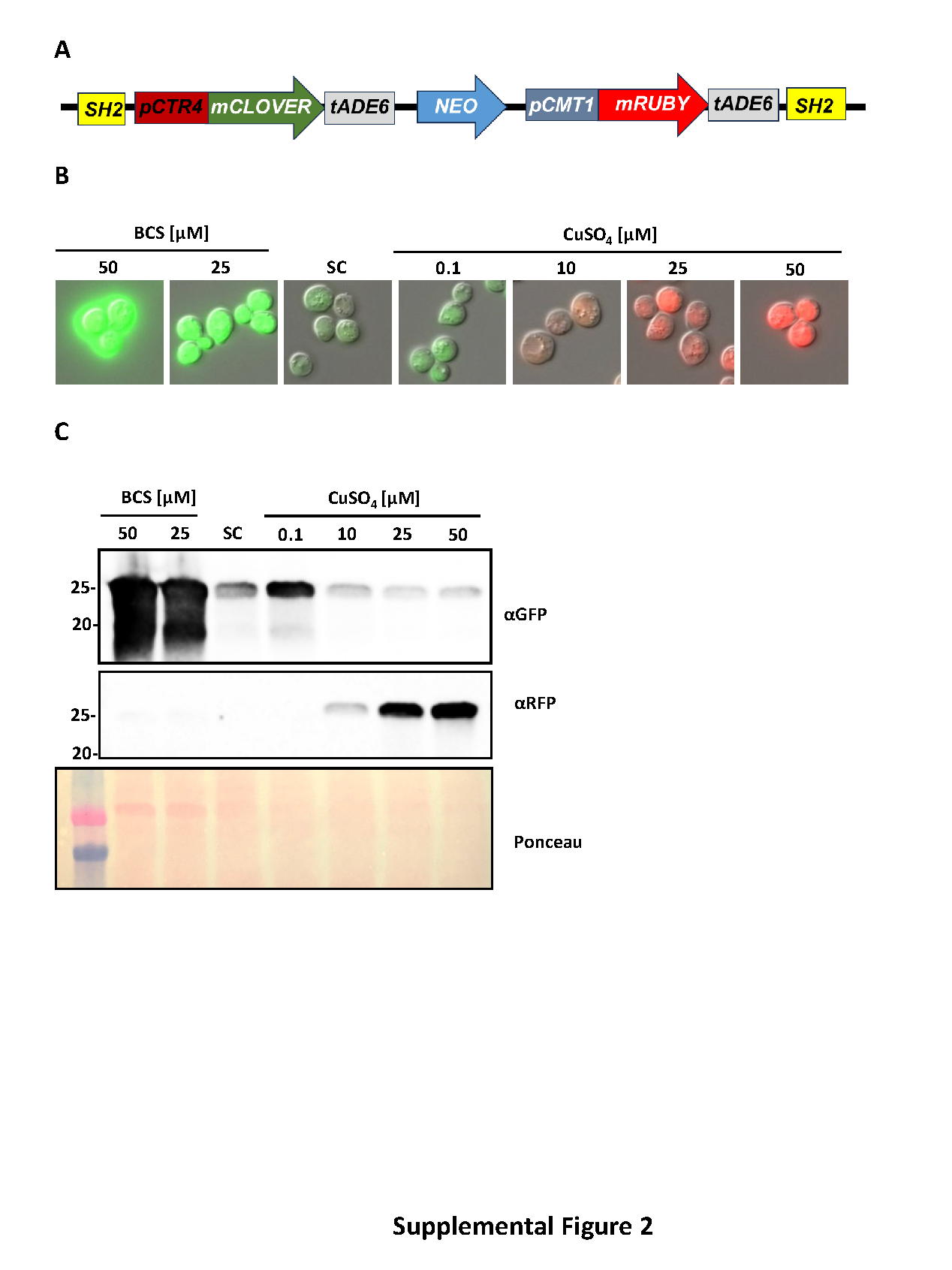

### Supp Fig 3

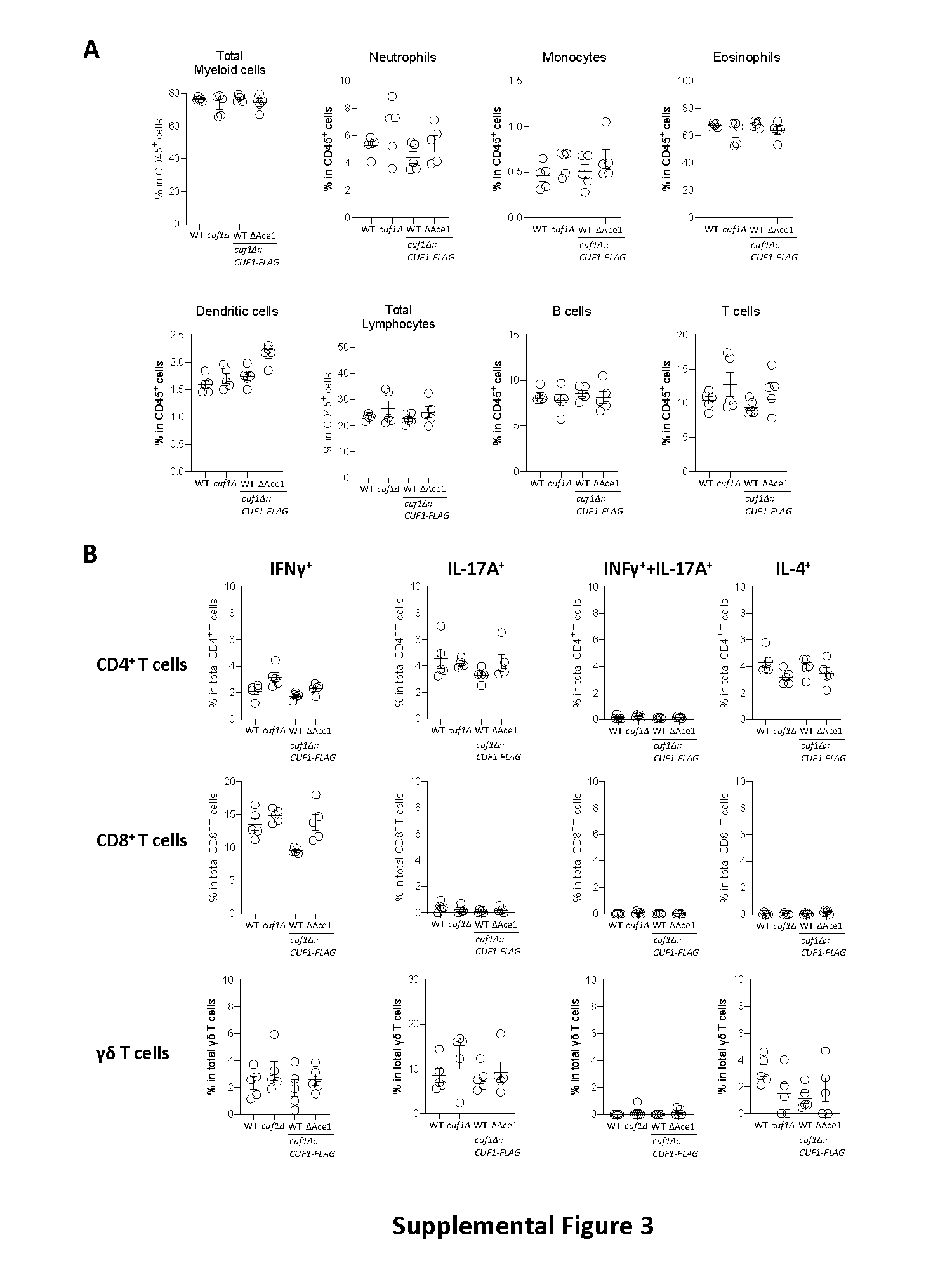

### Supp Tab5

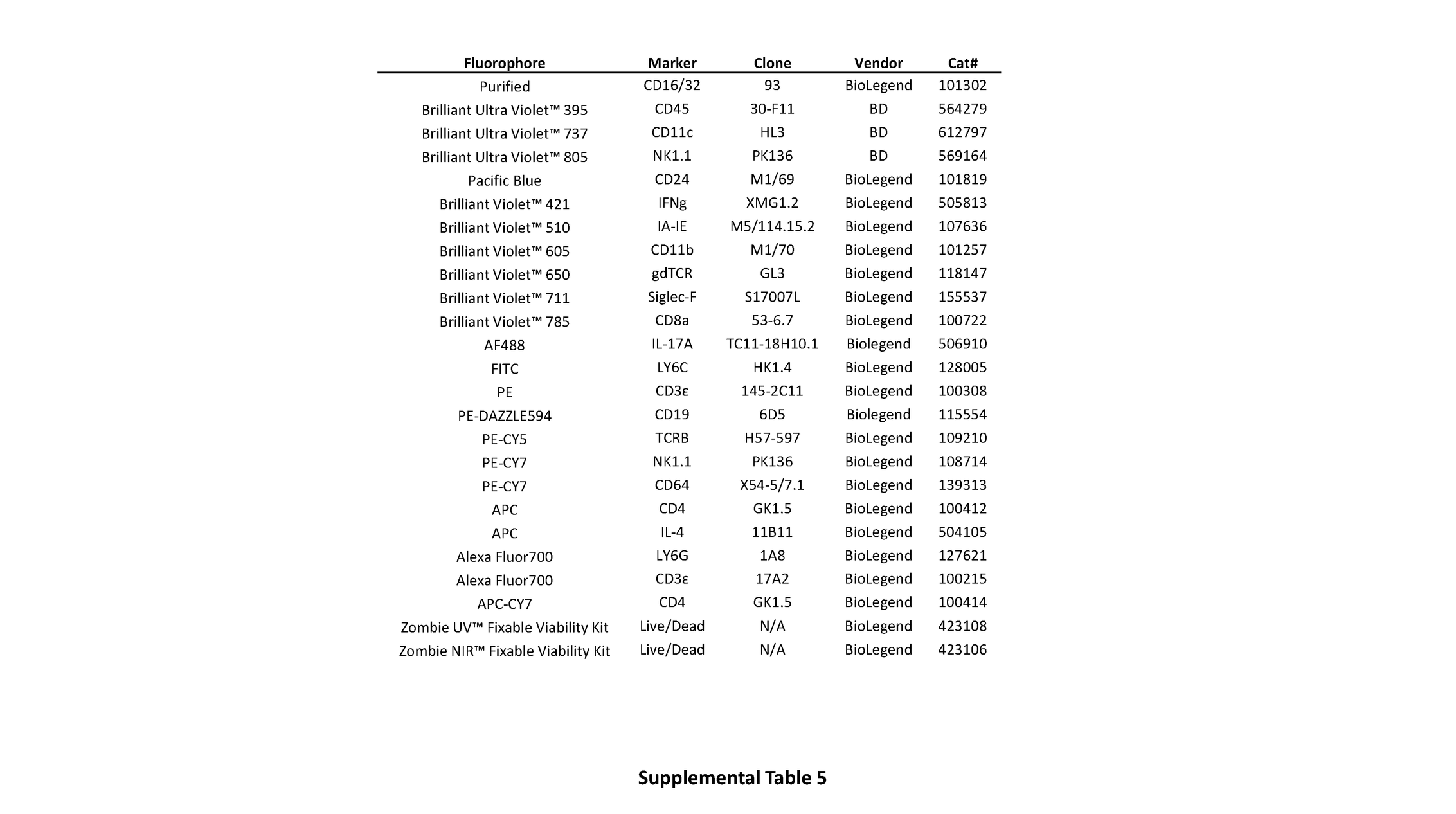

### Supp Tab 1

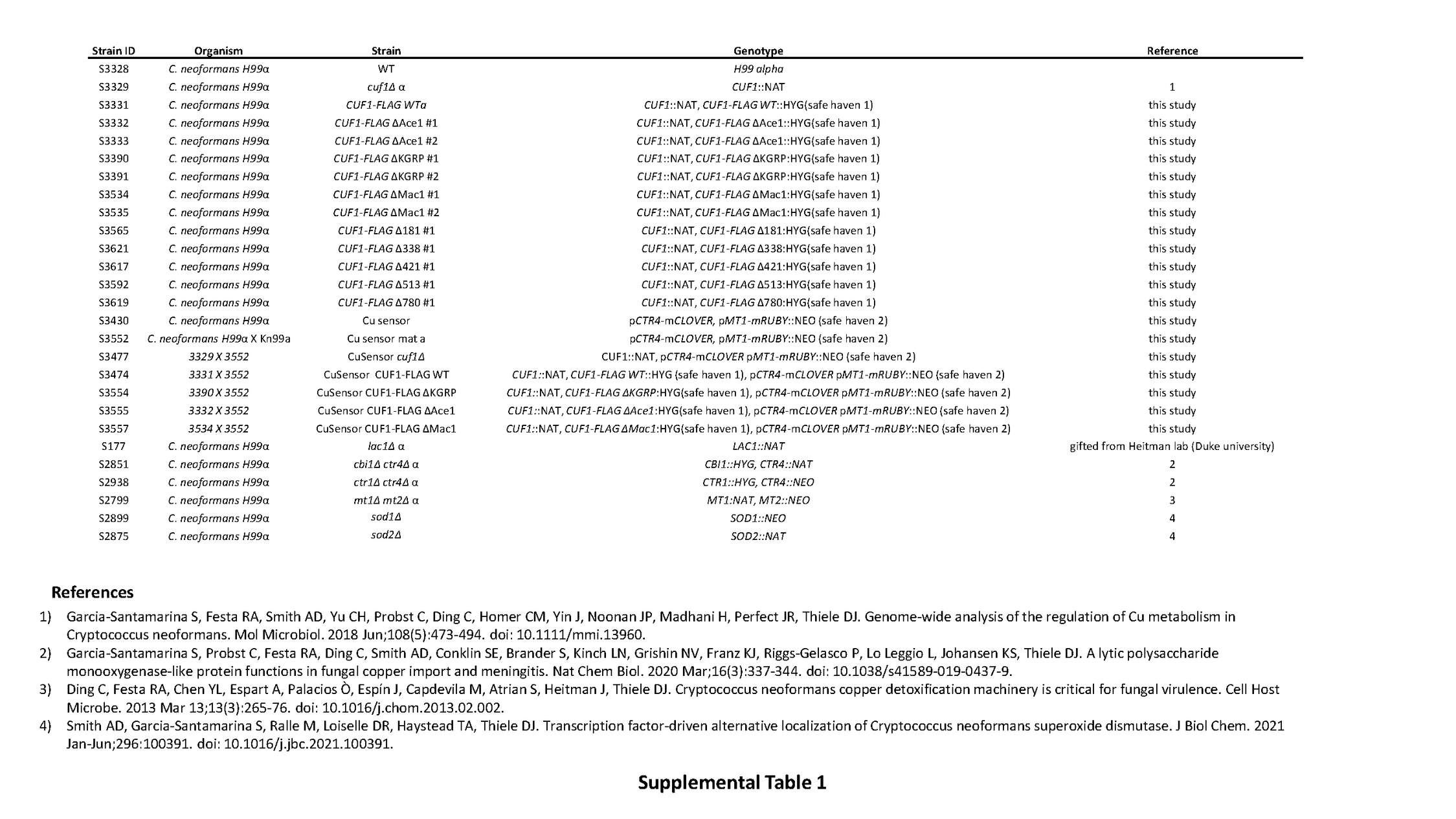

### Supp Tab 2

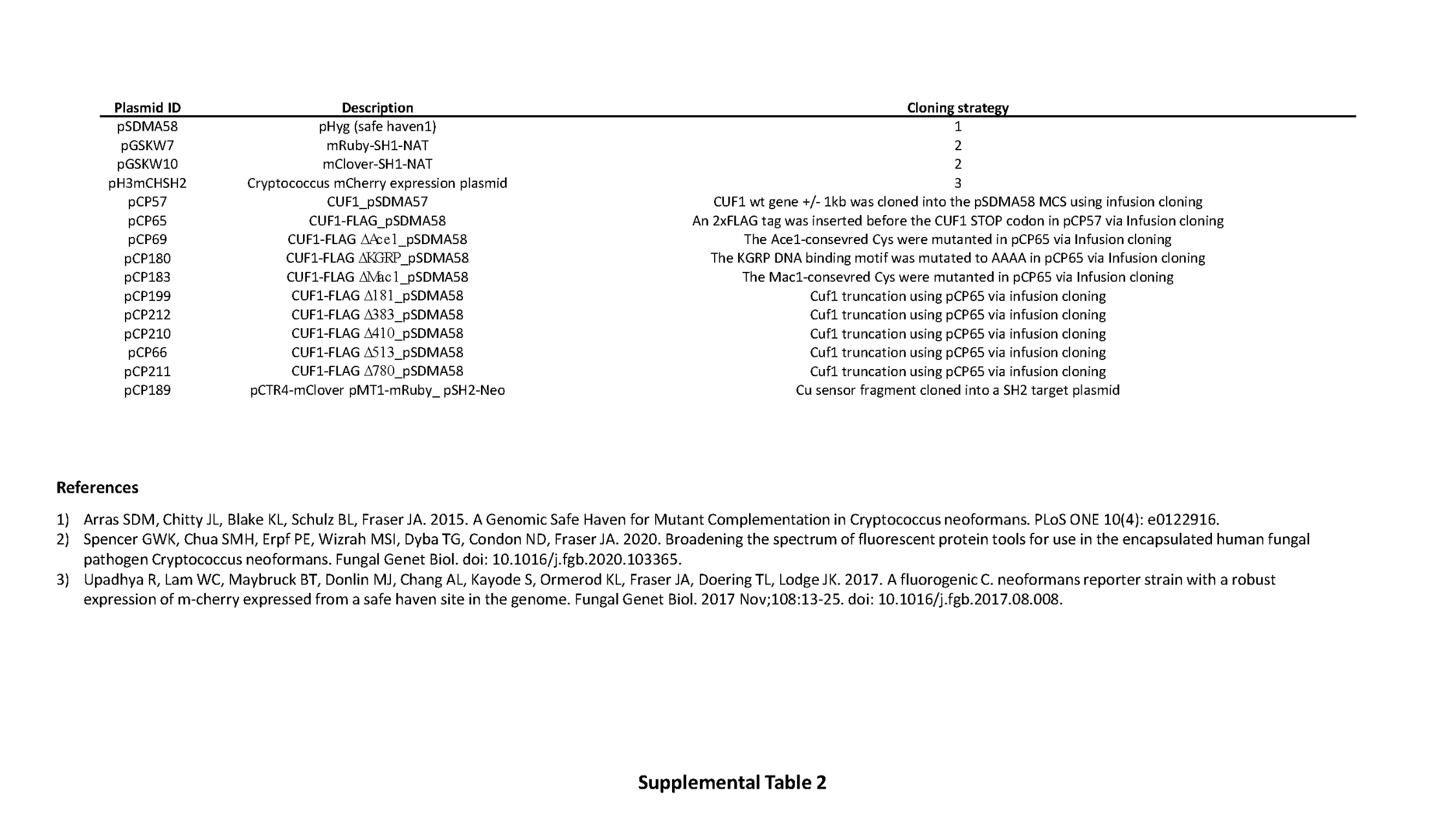

### Supp Tab 3

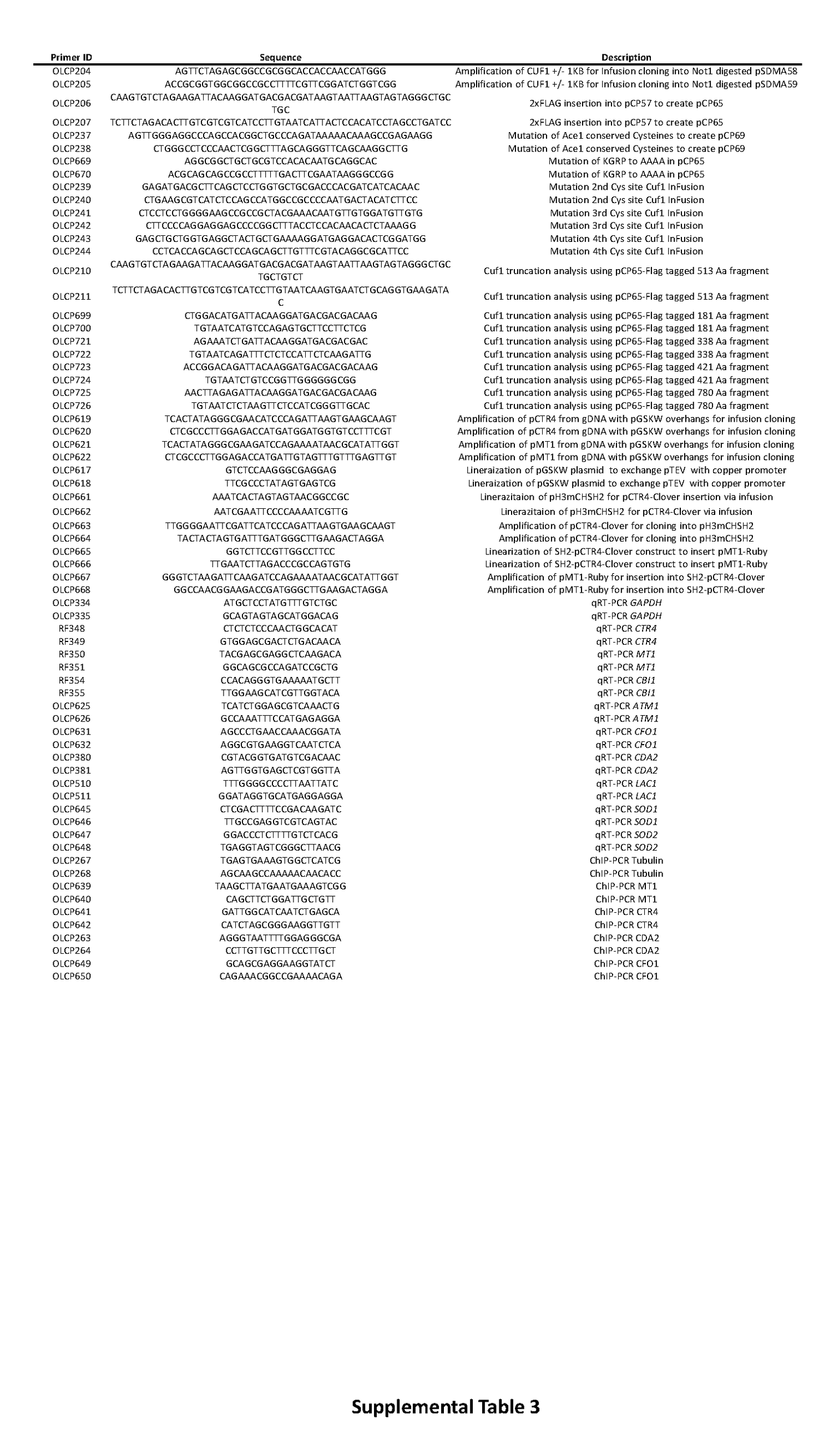

### Supp Tab 4

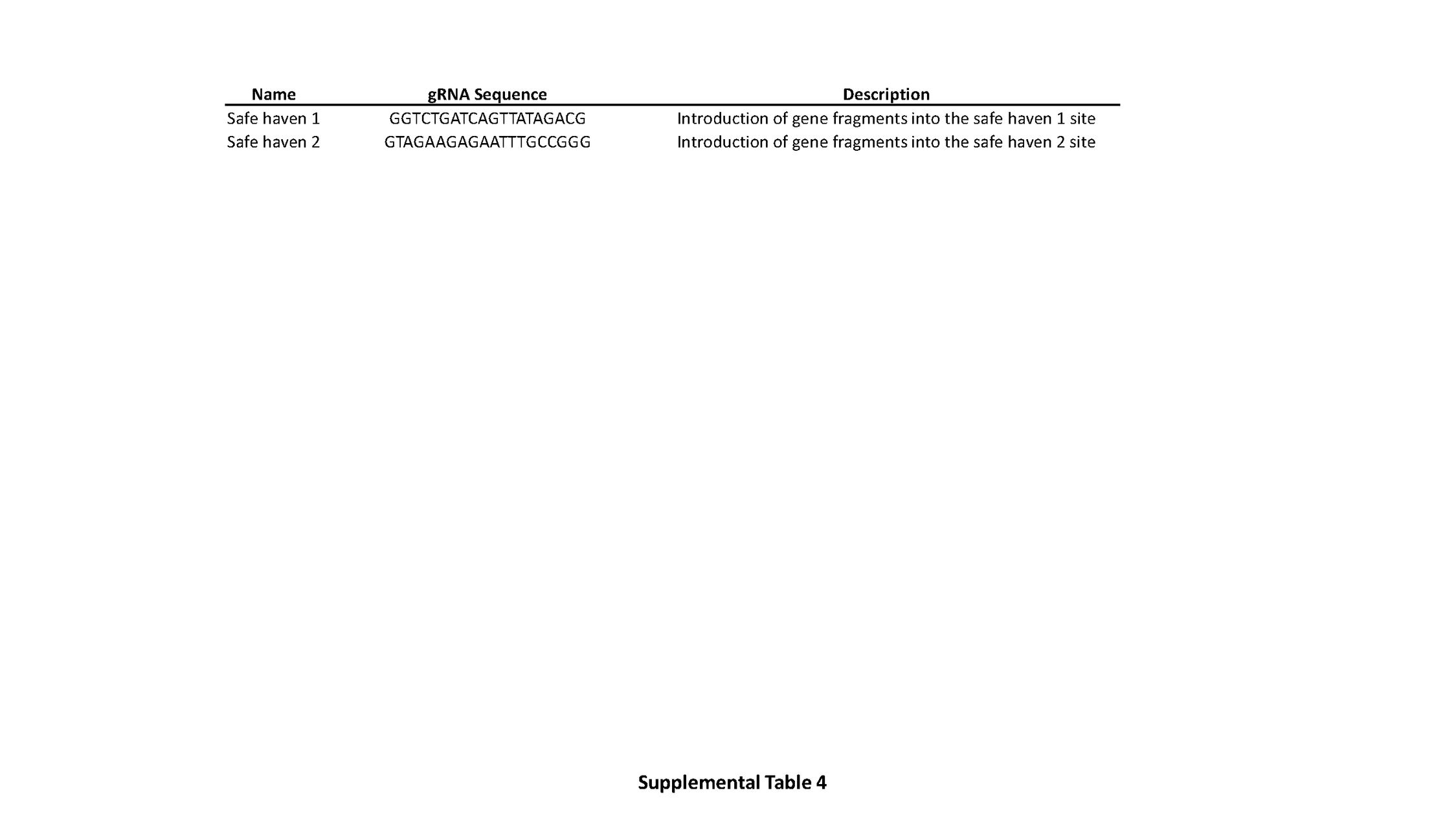
