## Supplementary material for "A cysteine-rich domain of the *Cryptococcus neoformans* Cuf1 transcription factor is required for high copper stress sensing and fungal virulence": Figure Legends

**SUPPLEMENTAL**

**Supplemental Figure S1:**

**(A-B)** Schematic representation of all generated Cuf1-FLAG motif mutants (A) and truncations (B). The first and last residues of each domain are indicated. The N-terminal domain is shown in light grey; the C-terminal domain is shown in dark grey. The N-terminal DNA binding motif, which contains the conserved KGRP DNA binding motif, is highlighted in yellow. The Ace1-like cysteine-rich region is indicated via blue box, and the Mac1-like regions are indicated by an orange box. The sequence of the identified cysteine-rich regions is indicated for each region with the *Cn*Cuf1-unique Cys-rich sequence located adjacent to the 2^nd^ Mac1-like Cys motif is highlighted in turquoise. Cysteine (Cys, C) to alanine (Ala, A) are indicated for each site mutant and are highlighted in red.

**Supplemental Figure S2:**

**(A)** Schematic representation of the p*CTR4*-*mCLOVER* and p*CMT1*-m*RUBY* containing Cu sensor allele inserted into the *Cn* safe haven 2 genomic locus. **(B-C)** Microscopic (B) and western blot mediated (C) validation of Cu-stress dependent expression of *mCLOVER* and *mRUBY* in the generated WT Cu sensor strain. The Cu sensor strain was inoculated to OD600 of 0.1 and cultured for 24 h in SC supplemented with indicated concentrations of BCS or CuSO_4_. (B) Microscopic validation was performed using the GFP and Texas Red channel of a Zeis Axio Imager M2. Image analysis was performed using ImageJ/Fiji. (B) Western blot validation was performed using TCA extracts of growth cultures. 20 µg total protein was loaded per lane. mClover was detected using a monoclonal anti-GFP antibody (mouse, αGFP) and mRuby was detected using a polyclonal anti-RFP antibody (rabbit, αRFP). A Ponceau staining was performed to control for protein loading and transfer.

**Supplemental Figure S3:**

Flow cytometry analysis of changes in various immune cell populations **(A)** and T-cell activation **(B)**. Female A/J mice were infected by inhalation with 10^5^ cells of the indicated strains (5 mice/strain). Mice were sacrificed 14 days postinfection and their lungs harvested. Relative percentages of immune cell subtypes are shown. Data were plotted via GraphPad Prism.

**Supplemental Table S1: Strains used in this study.**

**Supplemental Table S2: Plasmids used in this study.**

**Supplemental Table S3: Oligonucleotides used in this study.**

**Supplemental Table S4: Guide RNA used for CRISPR/Cas9 mediated transformation**

**Supplemental Table S5: Antibodies used for flow cytometry analysis**
